# Discovery of putative Golgi S-Adenosyl methionine transporters reveals the importance of plant cell wall polysaccharide methylation

**DOI:** 10.1101/2021.07.06.451061

**Authors:** Henry Temple, Pyae Phyo, Weibing Yang, Jan J. Lyczakowski, Alberto Echevarría-Poza, Igor Yakunin, Juan Pablo Parra-Rojas, Oliver M. Terrett, Susana Saez-Aguayo, Ray Dupree, Ariel Orellana, Mei Hong, Paul Dupree

## Abstract

Polysaccharide methylation, especially that of pectin, is a common and important feature of land plant cell walls. Polysaccharide methylation takes place in the Golgi apparatus and therefore relies on the import of S-adenosyl methionine (SAM) from the cytosol into the Golgi. However, to date, no Golgi SAM transporter has been identified in plants. In this work, we studied major facilitator superfamily members in Arabidopsis that we identified as putative Golgi SAM transporters (GoSAMTs). Knock-out of the two most highly expressed GoSAMTs led to a strong reduction in Golgi-synthesised polysaccharide methylation. Furthermore, solid-state NMR experiments revealed that reduced methylation changed cell wall polysaccharide conformations, interactions and mobilities. Notably, the NMR revealed the existence of pectin ‘egg-box’ structures in intact cell walls, and showed that their formation is enhanced by reduced methyl-esterification. These changes in wall architecture were linked to substantial growth and developmental phenotypes. In particular, anisotropic growth was strongly impaired in the double mutant. The identification of putative transporters that import SAM into the Golgi lumen in plants provides new insights into the paramount importance of polysaccharide methylation for plant cell wall structure and function.

---

All plant cells are enclosed by a network of cell wall polysaccharides. This network must be strong enough to resist internal pressures yet remain flexible to permit cell growth. The balance between these attributes is governed by the properties of the cell wall, which are determined by the structure, chemistry and the interactions of its constituent polysaccharides.

With the exceptions of cellulose and callose, cell wall polysaccharides are synthesised and modified in the Golgi apparatus, where various enzymes catalyse the transfer of glycosyl, acetyl, and methyl moieties onto glycan acceptors from a range of donor substrate molecules. Many of these donor substrates need to cross the Golgi membrane barrier in order to reach the lumen of the organelle. To achieve this, the Golgi membrane contains several types of transporter proteins that import essential metabolites, such as nucleotide sugars, acetylation donors, and the methylation donor *S*-adenosyl methionine (SAM) ^1,2^.

Various residues can be methylated in plant cell wall polysaccharides: glucuronic acid (GlcA) of xylan and arabinogalactan proteins ^3,4^, various sugars in the pectin rhamnogalacturonan II (RG-II) ^5^, and galacturonic acid (GalA) in the pectin homogalacturonan (HG), the most abundantly methylated polysaccharide ^6^. The degree of HG methylation is considered a major factor influencing the capacity of cells to expand ^7–9^. The ‘egg-box’ model describes the capacity of HG with a low degree of methyl-esterification (DM), which is negatively charged, to form Ca^+2^-mediated ionic cross-links to other HG molecules, a process that may lead to cell wall stiffening and restriction of cell growth ^10^. Although the ‘egg-box’ model is widely accepted, there is no direct evidence supporting the existence or extent of these structures in intact plant cell walls. On the other hand, low-DM HG serves as a substrate for polygalacturonase enzymes (PGAses) present *in muro*, causing HG degradation that has been associated with cell wall loosening and cell expansion ^11–13^. The degree of HG methylation *in muro* is tightly regulated by the interplay between pectin methyl esterases (PMEs), enzymes able to remove methyl groups from HG, and their inhibitors (PME inhibitors; PMEIs) ^7^.

Biochemistry and immunogold experiments have shown that pectin methylation takes place in the Golgi apparatus ^14,15^, but little is known about the different activities that facilitate this process. Recently, QUA2/TSD2 was shown to methylate HG using SAM as donor substrate *in vitro*, becoming the first candidate pectin methyl transferase (PMT) to be biochemically characterised^16^. Mutations in this enzyme cause severe growth and cell adhesion phenotypes ^17,18^. However, there was no evident decrease in the absolute degree of pectin methylation in mutants of QUA2 or the related DUF248 family protein QUA3 ^16,19^. Genetic redundancy or specialisation of activity of the large number of related DUF248 candidate PMTs to restricted pectin domains makes the study of the general importance of pectin methylation difficult. The CGR2 and CGR3 proteins comprise a distinct family of two putative PMTs. The *cgr2 cgr3* mutant exhibits marked reduction in HG methylation and presents severe growth phenotypes ^20^. However, *in* vitro assays using microsomes of CGR2 and CGR3 overexpression lines only led to a slight increase in methyltransferase activity using SAM and homogalacturonan as substrate. In addition, the *cgr2 cgr3* mutant showed no significant decrease in activity when compared to WT; therefore, the actual substrate of these enzymes *in planta* is unclear.

Importantly, all pectin methylation depends on the presence of SAM in the Golgi lumen; therefore, methylation is underpinned by the presence of Golgi SAM transporters (**Fig1A**). While Golgi SAM transport has been detected in plants ^21^, to date, no candidate for a plant Golgi SAM transporter has been studied. In this work, we identified three members of the major facilitator superfamily (MFS) as putative Golgi-localised SAM transporters. Knocking-out the two most abundantly expressed members leads to reduced methylation of Golgi synthesised polysaccharides as well as to severe growth and developmental phenotypes. Using cell wall biochemistry and solid-state NMR (ssNMR) experiments, combined with genetic tools, we established that these putative transporters play a major role in the methylation of pectin and xylan, and that reduced GoSAMT activity leads to important changes in the molecular architecture of the plant cell wall.

**Figure 1.**
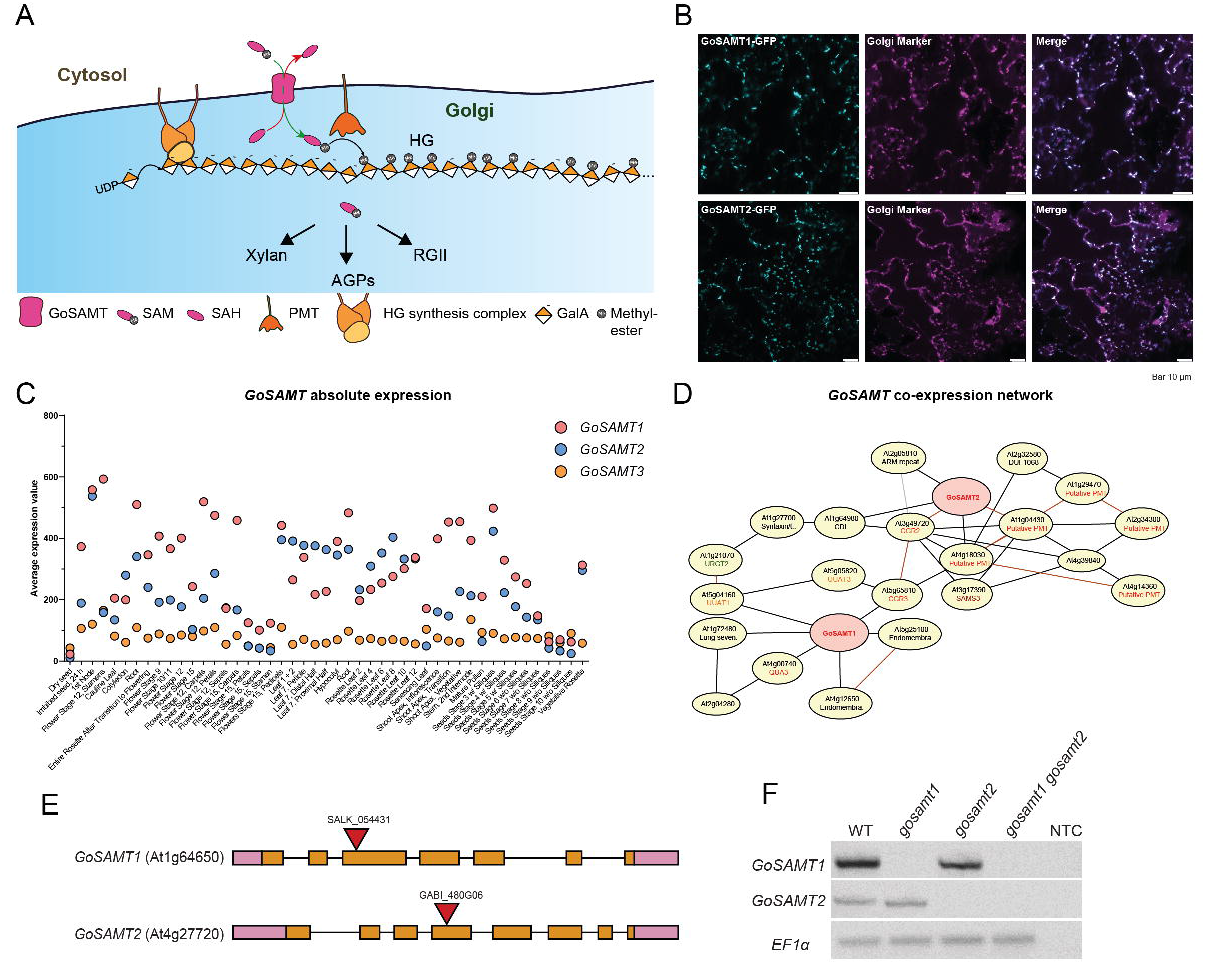
Identification of putative Golgi SAM Transporters. (**A**) Scheme describing the importance of GoSAMTs in the process of polysaccharide methylation in the Golgi apparatus. (**B**) Subcellular localisation of GoSAMT1-GFP and GoSAMT2-GFP expressed under their endogenous promoters in stable lines of Arabidopsis plants. (**C**) and (**D**) Arabidopsis expression and co-expression network data from the eFP browser and Atted-II respectively, data show *GoSAMT1* and *GoSAMT2* are highly and ubiquitously expressed in Arabidopsis and they cluster in the same co-expression network with many putative pectin methylation genes. (**E**) Schematic of *GoSAMT1* and *GoSAMT2* genes including the sites of T-DNA insertions. (**F**) RT-PCR experiments with primers to amplify *GoSAMT1* and *GoSAMT2* genes, using cDNA template from WT, *gosamt1*, *gosamt2* and *gosamt1 gosamt2* mutants. NTC, no template control.

## Results

### Identification of putative Golgi SAM transporters in plants

To identify candidate Golgi SAM transporters, we searched for proteins in the Arabidopsis proteome with characteristics consistent with this activity. We examined multipass transmembrane proteins shown to be abundant in the Golgi based on our previous proteomic experiments of Arabidopsis root callus ^22,23^. Amongst candidates for Golgi SAM transporters, we found two members of the major facilitator superfamily 5 (MFS_5), AT1G64650 and AT4G27720, ranked in the top 50 most abundant proteins of this organelle ^23^. MFS proteins are ubiquitously distributed across organisms, playing roles in the transport of a broad spectrum of ions and solutes across membranes ^24^. Some MFS_5 proteins, such as *Homo sapiens* MFSD5 and *Chlamydomonas* MFS_5, have been identified as members of a molybdate transporter family, MOT2 ^25^. However, there is no biochemical or subcellular localisation evidence that these are plasma membrane molybdate transporters ^25^. On the other hand, an orthologue in *C. elegans* (SAMT1) has been proposed to act as a Golgi SAM transporter due to the fact that its mutation confers resistance to Tectonin2—a toxin that relies on the presence of methylated extracellular glycans ^26^.

There are three closely related MFS_5 proteins in Arabidopsis and given the data below and their resemblance to *C. elegans SAMT1* (**Fig.S1A**), we named these proteins <u>Go</u>lgi <u>S</u>-<u>A</u>denosyl <u>M</u>ethionine <u>T</u>ransporters 1-3 (AT1G64650 (GoSAMT1), AT4G27720 (GoSAMT2) and AT3G49310 (GoSAMT3)). There are related proteins in many eukaryotes and in all Viridiplantae species analysed, from green algae to embryophytes (**Fig.S1C**). Topology analysis using the tool TOPCONS (http://topcons.cbr.su.se/) predicted that *C. elegans SAMT1* and the three Arabidopsis members have 13 hydrophobic domains (**Fig.S1B**), of which the first may correspond to a cleaved signal peptide followed by 12 transmembrane helices. To confirm Golgi localisation of these proteins, we generated stable transgenic Arabidopsis lines of *GoSAMT1-GFP* and *GoSAMT2-GFP*, expressed under their endogenous promoters. Both proteins co-localised with the Golgi marker mannosidase-I (Man-I) (**Fig.1B**), indicating that they are localised in the Golgi apparatus—consistent with the previous proteomic data ^22,23^.

To investigate whether GoSAMTs play a role in polysaccharide methylation, we first looked at co-expressed genes in the ATTED-II database (http://atted.jp/) ^27^. The analysis showed that *GoSAMT1* and *GoSAMT2* cluster in a co-expression network that contains predominantly pectin synthesis and methylation genes, suggesting the encoded proteins could be playing a role in pectin methylation (**Fig.1D**). Expression data from the eFP Browser (https://bar.utoronto.ca/efp/cgi-bin/efpWeb.cgi) ^28^ shows that *GoSAMT1* and *GoSAMT2* are widely expressed across different tissues and developmental stages in Arabidopsis, while the expression of *GoSAMT3* is considerably lower (**Fig.1C**), in line with our previous quantitative Golgi proteomic experiments ^23^. Therefore, we focused our search for genetic evidence for a role of these MFS_5 proteins by studying *gosamt1* and *gosamt2* knock-out mutants. We identified and isolated homozygous T-DNA insertion mutants (**Fig.1E**), and we generated a double *gosamt1 gosamt2* mutant, confirming the lack of transcripts in the mutant plants by RT-PCR experiments (**Fig.1F**).

### Xylan and pectin methylation is affected in the *gosamt1 gosamt2* mutant

In contrast to mutations in specific polysaccharide methyltransferases, we reasoned that mutations in *GoSAMT* genes ought to affect polysaccharide methylation in a global manner. We began by analysing cell wall xylan GlcA methylation. We performed GH11 endoxylanase digestion of xylan from alcohol insoluble residues (AIR) of basal inflorescence stems, and the percentage of GlcA methylation was evaluated by relative quantitation of the released oligosaccharides Xyl_4_GlcA and Xyl_4_^Me^GlcA using capillary electrophoresis ^29^. Xylan GlcA of the *gosamt1* mutant was less methylated than xylan of WT plants (**Fig.2A**). Although the xylan methylation in *gosamt2* mutants was not different to that of WT plants, the GlcA methylation of *gosamt1 gosamt2* double mutant xylan was reduced by two thirds, when compared to WT, indicating that both putative transporters play a role in secondary cell wall xylan methylation. Molecular complementation experiments restored xylan GlcA methylation to WT levels when we expressed *_pro_GoSAMT1:GoSAMT1-GFP* in *gosamt1 gosamt2* mutants (**Fig.S2D**). Although we did not observe restoration when we expressed *_pro_GoSAMT2:GoSAMT2-GFP*, this may be due to technical reasons such as the construct missing regulatory elements for expression in secondary cell wall synthesis. The reduction of xylan GlcA methylation in *gosamt1* single and *gosamt1 gosamt2* double mutant plants is clear evidence that these MFS_5 proteins participate in xylan methylation.

**Figure 2.**
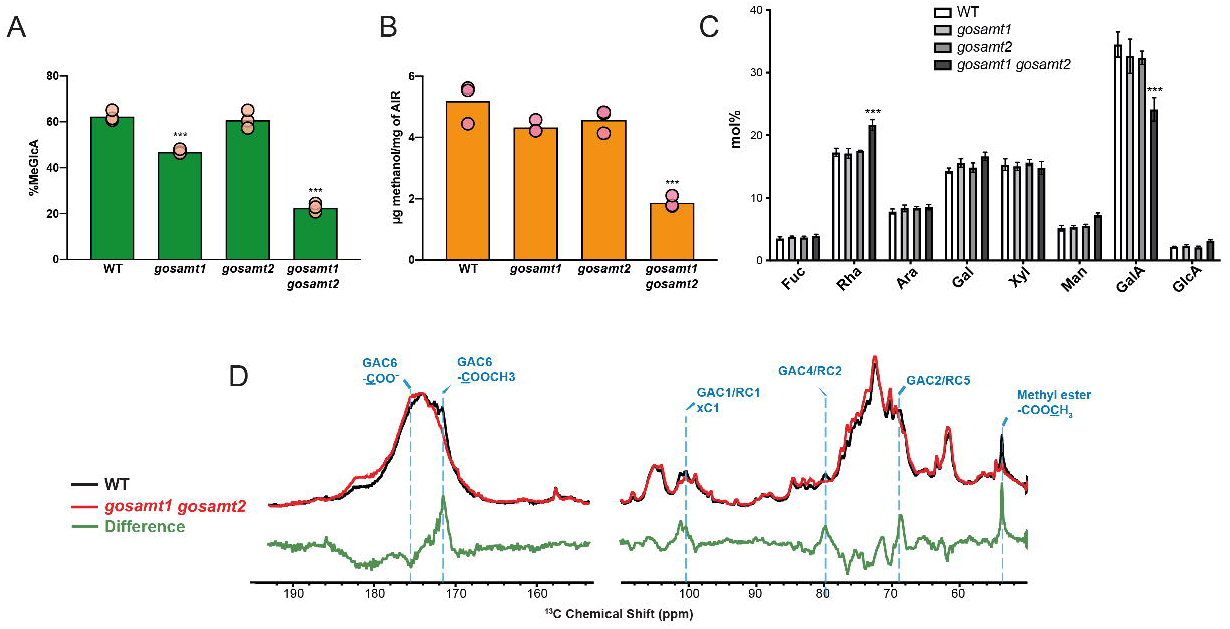
*GoSAMT* mutants show reduced polysaccharide methylation. (**A**) Ratio of released Xyl_4_GlcA and Xyl_4_^Me^GlcA products after endoxylanase treatment of basal stem AIR, followed by capillary electrophoresis separation of WT, *gosamt1*, *gosamt2* and *gosamt1 gosamt2* mutants. Values correspond to the mean of 3 biological replicates. Asterisks indicate significant differences between mutants and the WT defined by one-way ANOVA followed by Bonferroni’s multiple comparison test: ***, P < 0.001. (**B**) Quantification of methanol release after saponification of leaf AIR of WT, *gosamt1*, *gosamt2* and *gosamt2 gosamt2*. Values correspond to the mean of 3 biological replicates. Asterisks indicate significant differences between mutants and the WT defined by one-way ANOVA followed by Bonferroni’s multiple comparison test***, P < 0.001. (**C**) HPAEC-PAD monosaccharide composition analysis of leaf AIR of WT (white), *gosamt1*, *gosamt2* (light and dark grey respectively) and *gosamt1 gosamt2* (black), values correspond to the mean and ± SEM of 4 biological replicates. Asterisks indicate significant differences between mutants and the WT defined by two-way ANOVA followed by Tukey’s multiple comparison test: *, P < 0.05; **, P < 0.01; ***, P < 0.001. Ara, arabinose; Fuc, fucose; Gal, galactose; GalA, galacturonic acid, GlcA, glucuronic acid; Man, mannose; Rha, rhamnose; Xyl, xylose (**D**) 20s DP 1D ^13^C ssNMR spectrum of WT (black) and *gosamt1 gosamt2* (red) of never-dried ^13^C enriched leaves. The mutant shows reduced GalA signals, as well as a marked reduction in COO<u>**C**</u>H_3_ and <u>**C**</u>OOCH_3_, whereas it shows increased <u>**C**</u>OO^−^ signals.

We next investigated whether the loss of GoSAMTs affects pectin methyl-esterification. Methyl-esterified HG is the main source of methanol released after saponification, so we performed a methanol release assay on cell wall material obtained from rosette leaves of adult plants. Compared to WT plants, both *gosamt1* and *gosamt2* single mutants released less methanol, by 17% and 12% respectively. Remarkably, in the case of the *gosamt1 gosamt2* mutant, we observed a severe reduction in methanol release of about two thirds of WT levels (**Fig.2B**), similar to the scale of reduction seen for xylan GlcA methylation. The methanol content in the *gosamt1 gosamt2* mutant was restored to single mutant values in the respective lines complemented with each gene (**Fig.S2E**), further supporting the finding that polysaccharide methylation is dependent on both GoSAMT1 and GoSAMT2.

To investigate if changes in polysaccharide methylation affect the sugar composition of cell walls, we performed monosaccharide composition analysis of leaf AIR. The *gosamt1 gosamt2* plants have a decrease of 25% in relative GalA content compared to WT (**Fig.2C**). Despite this decrease in GalA content in *gosamt1 gosamt2*, the substantially greater decrease in methanol content suggests that the remaining HG in *gosamt1 gosamt2* is less methyl-esterified than in the WT. To investigate this further, we performed quantitative 1D solid-state NMR experiments on never-dried, ^13^C labelled leaves of WT and *gosamt1 gosamt2* mutant plants. The ^13^C NMR spectra, using direct polarisation (DP) and a recycle delay of 20 s, gives an accurate report of the relative proportions of different polysaccharides and functional groups present in cell walls ^30,31^. There were few changes between WT and mutant, except in pectin-related signals. There was a marked decrease in the mutant in the 53.7-ppm peak, which corresponds to the methyl ester group on the GalA COO<u>C</u>H_3_ (**Fig.2D**). Furthermore, the peak of carbon 6 for methyl esterified GalA <u>C</u>OOCH_3_ (GAC6) was clearly reduced, whereas the unmethylated galacturonate <u>C</u>OO^−^ signal was notably higher (**Fig.2D**). Although not fully resolved from other components in the cell wall in the 1D spectrum, the GalA ^13^C peaks at 101 ppm (GAC1), 80 ppm (GAC4), and 69 ppm (GAC2), showed weaker signals in the mutant spectrum compared to the WT spectrum (**Fig.2D**). These NMR results therefore support the results of our biochemical analysis, indicating HG content is somewhat reduced in the *gosamt1 gosamt2* mutant, and that the HG has substantially lower methyl-esterification.

### ssNMR experiments reveal important changes in the cell wall molecular architecture of the *gosamt1 gosamt2* mutant

To investigate how the HG esterification state influences pectin conformations and inter-molecular interactions, we next studied ^13^C-labelled, never-dried leaf material using 2D ssNMR experiments. We first investigated whether the conformations of the cell wall polymers are affected in the *gosamt1 gosamt2* mutant. Short mixing time cross-polarisation proton-driven spin diffusion (CP-PDSD) experiments were used to study the cell wall composition and chemical shifts of the main carbohydrate components (**Fig.3**). At a mixing time of 100 ms, where cross-peaks are largely intra-residue, the WT and mutant PDSD spectra showed similar ^13^C chemical shifts and linewidths. Thus, any intensity differences indicate the concentration changes of polysaccharides in the *gosamt1 gosamt2* mutant. We observed that different cellulose environment cross-peaks, for example internal and surface cellulose cross-peak signals (iC4-iC6 89 ppm, 65 ppm; sC4-sC6 84 ppm, 62.5 ppm), displayed similar intensities between the WT and *gosamt1 gosamt2* cell walls. This indicates that the cellulose fibril nanostructure and cellulose content are unchanged in this mutant. While cellulose signals displayed similar intensities between WT and *gosamt1 gosamt2*, pectin polysaccharides signals were clearly reduced in the mutant, and they showed notable changes. HG ^13^C signals at 101 ppm (GAC1), 80 ppm (GAC4), and 68.6 ppm (GAC2) are 15-25% weaker in the mutant spectrum compared to the WT spectrum. In addition, GalA cross-sections, such as at 68.6 ppm (GAC2) (**Fig. 3B**), showed that the *gosamt1 gosamt2* mutant spectrum contains lower intensity for the 172 ppm peak of the methyl esterified carbon 6 GAC6 (O<u>C</u>OCH_3_). These differences provide further evidence for the lower degree of HG methyl-esterification in the mutant. Curiously, the *gosamt1 gosamt2* mutant spectrum displayed a higher intensity of the cellulose-bound xylan Xn4^2f^-Xn5^2f^ (82.2 ppm, 64.2 ppm) cross peak than the WT spectrum.

**Figure 3.**
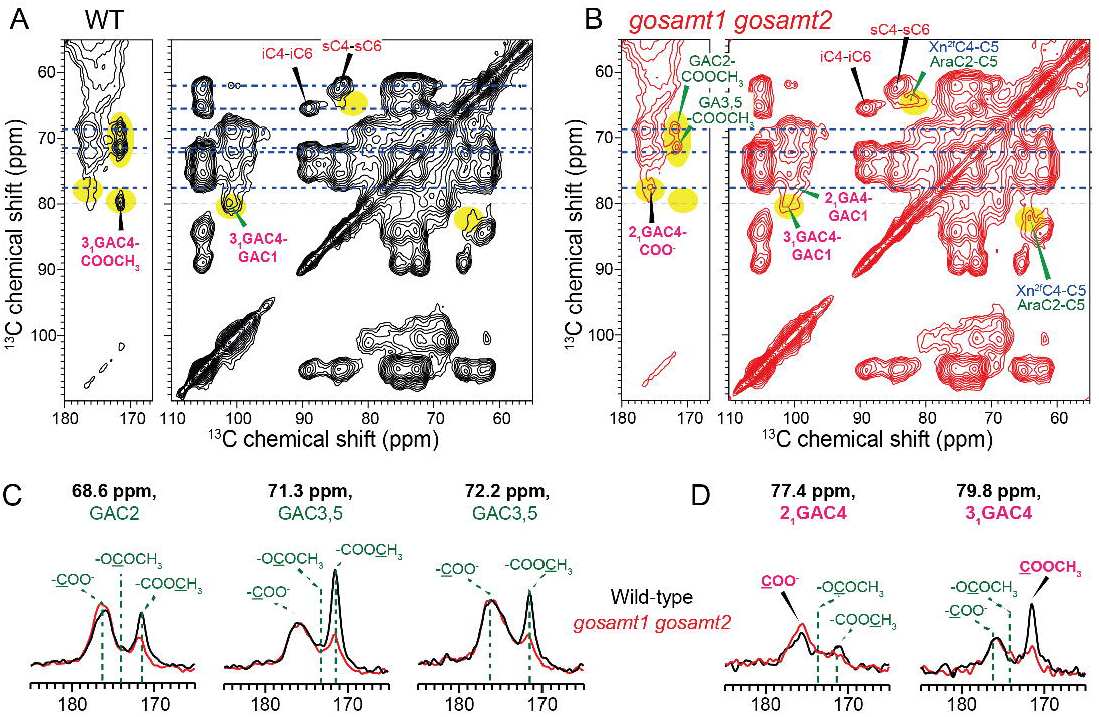
2D ssNMR experiments show marked reduction of pectin methyl-esterification and increased ‘egg-box’ structures. 100 ms 2D ^13^C CP-PDSD spectra of (**A**) WT and (**B**) *gosamt1 gosamt2* never-dried ^13^C enriched leaves. Highlighted regions (yellow) illustrate some of the differences between the two spectra. Pink annotations correspond to 2_1_ ‘egg-box’ signals. (**C**) Representative pectin cross sections. *gosamt1 gosamt2* displays much lower intensity for the 172 ppm <u>C</u>OOCH_3_ cross peak than WT but higher 176 ppm carboxyl peak intensity. The cross sections are scaled by the integrated area of each cross section between 0-200 ppm. (**D**) Cross sections at 77.4 ppm and 79.8 ppm, which contain the C4 signals of GalA in the calcium-crosslinked 2_1_ egg-box conformation and in the 3_1_ conformation, respectively. The mutant spectrum shows higher intensity for the 2_1_ conformation and lower intensity for the 3_1_ conformation compared to the WT (assigned in magenta). The corresponding positions in the 2D spectrum are shown in magenta. For clarity, only intra residue cross-peak signals from major wall components of interest are labelled.

Low-DM HG chains can form calcium-mediated dimers known as 2_1_ (2-fold screw) conformation ‘egg-box’ structures. 1D ^13^C NMR experiments have been used to monitor the HG conformational changes that occur when HG binds to Ca^+2^ *in vitro*. It was shown that the GAC4 79.8 ppm chemical shift of the 3_1_ conformation of HG changes to 77.4 ppm in the 2_1_ ‘egg-box’ conformation ^32^. Similar 1D ^13^C NMR experiments suggested the presence of this 2_1_ conformation in dried onion cell walls, however the congested area around the 77.4 ppm peak in the 1D spectra made this difficult to confirm. To investigate the existence and any changes in these ‘egg-box’ structures *in planta*, we studied GAC4 chemical shifts in our Arabidopsis, never-dried, samples. Indeed, chemical shifts corresponding to both HG conformations were detected. The 100 ms PDSD spectra revealed strong cross-peaks between the 2_1_ ‘egg-box’ conformation GAC4 (2_1_ GAC4) 77.4 ppm and the GAC6 (<u>C</u>OO^−^) 176 ppm, consistent with this conformation largely consisting of unesterified HG. Interestingly, these HG 2_1_ ‘egg-box’ signals were stronger in the *gosamt1 gosamt2* mutant in comparison to the WT. On the contrary, the cross-peak of the 3_1_ conformation GAC4 (3_1_GAC4) 79.8 ppm to the GAC6 methyl ester (O<u>C</u>OCH_3_) 172 ppm was stronger in the WT (**Fig. 3D**). Therefore, reduced pectin methyl-esterification in the mutants leads to the formation of more 2_1_ ‘egg-box’ HG structures.

We then investigated if the differences in the pectin structure of *gosamt1 gosamt2* walls have an impact on the mobility of the wall polysaccharides. We measured ^13^C-^1^H dipolar couplings using DIPSHIFT experiments. Quantitative DIPSHIFT spectra report the dynamics of ~85% of all matrix polysaccharides and also the majority of the signals for cellulose, which has the longest ^13^C T_1_ relaxation time among all wall polysaccharides ^33^. The experiment showed that many matrix polysaccharides display weaker dipolar dephasing, hence more mobility, in the *gosamt1 gosamt2* compared to WT samples (**Fig4**). These include, for example, the 82 ppm peak arising from arabinose C2/4 (AC2/4) and Xn4^2f^, and the 70.4 ppm peak arising from several Gal, GalA and Ara carbons ^33^. On the other hand, the pectin backbone regions, including GAC1 at 101 ppm and GAC4 at 80.2 ppm, show stronger dipolar dephasing for the mutant compared to WT. This indicates that the pectin backbone is more rigid in *gosamt1 gosamt2*.

**Figure 4.**
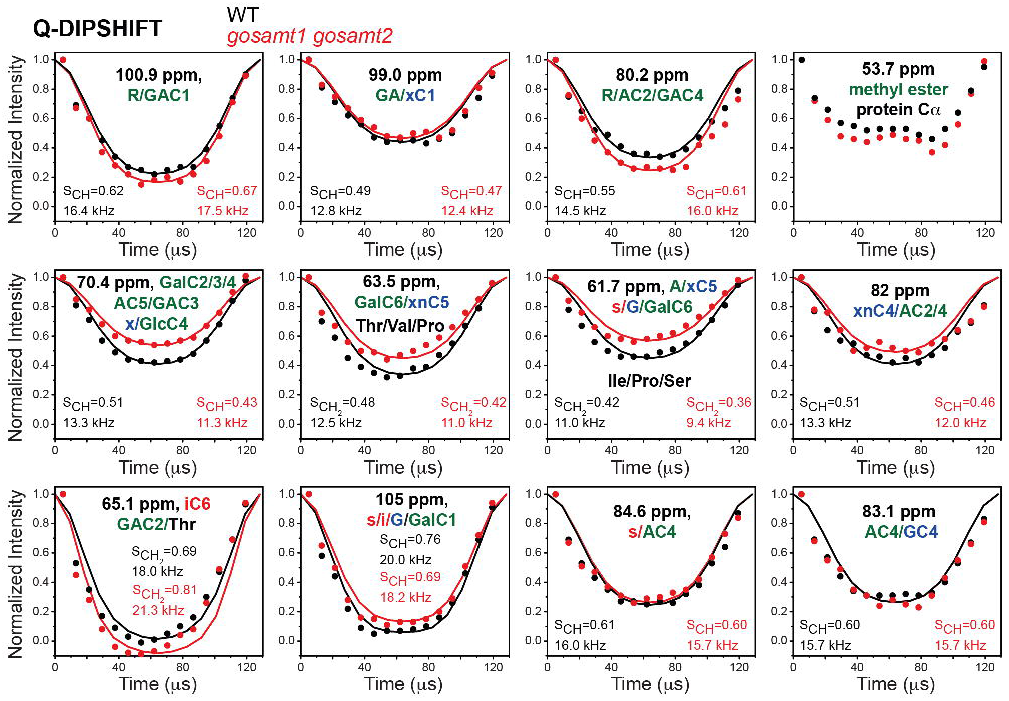
DIPSHIFT experiments show important changes in polysaccharide mobility. Quantitative DIPSHIFT curves of WT (black) and *gosamt1 gosamt2* (red) cell walls. Best-fit ^13^C-^1^H dipolar coupling values (scaled by FSLG) and S_CH_ values are given in each panel. The higher order parameters of the 100.9 ppm peak and the 80.2 ppm peak in *gosamt1 gosamt2* indicate that the pectin backbones are more rigid in the mutant cell wall than in the WT, while pectin side branches are more dynamic in the mutant.

To further probe pectin backbone immobilisation, we measured CP-DIPSHIFT spectra, which preferentially report the mobility of the more rigid polysaccharides, including cellulose and ~40% of matrix polysaccharides, an estimation based on the pectin intensity differences between 1D ^13^C CP and quantitative spectra ^12^. Cellulose-dominant peaks at 89 ppm, 65 ppm, and 105 ppm and the predominantly xyloglucan (XyG) peaks such as the 70.3 ppm peak show similar dipolar couplings between the mutant and WT cell walls, implying that cellulose and XyG are similarly rigid in the two cell walls (**Fig.S3**). On the contrary, the regions of the HG backbone signals at 101 ppm, 99.0 ppm, and 79.8 ppm show stronger dipolar couplings in the mutant than in the WT cell wall, indicating less mobility for this relatively immobile fraction of the wall. Moreover, *gosamt1 gosamt2* mutant cell walls also show stronger dipolar coupling for the 53.7 ppm peak. Although this peak overlaps with the methyl ester and protein Cα sites, the exclusive protein Cα peak at 56.2 ppm has identical dipolar couplings between the two cell walls; thus, the coupling increase at 53.7 ppm indicates that the mutant cell wall has more rigid HG methyl ester groups than the WT. These differences are further evidence supporting *gosamt1 gosamt2* plants have more rigid pectic polysaccharides.

To evaluate how a low level of pectin methyl-esterification affects polysaccharide intermolecular interactions, we measured 2D ^13^C-^13^C correlation spectra using a long PDSD mixing time of 1.5 s (**Fig. 5**). The 2D spectra showed important differences between WT and *gosamt1 gosamt2*. New signals were detected in *gosamt1 gosamt2* at 95.7 and 97.7 ppm. We attribute these to GAC1 reducing end (GAC1^re^) in their alpha and beta configuration, respectively. Remarkably, we saw clear cross-peaks among αGAC1^re^ and 2_1_ HG signals (95.7, 77.4 ppm; 95.7, 176 ppm) and absence of cross peak to COO<u>**C**</u>H_3_ (95.7, 54 ppm) or to <u>**C**</u>OOCH_3_ (95.7, 172 ppm) (**Fig. 5A**), suggesting *gosamt1 gosamt2* has short and abundant ‘egg-box’ structures. These cross-peaks are absent in the spectra of WT walls. Additionally, cellulose-pectin cross-peaks were resolved in the 2D spectra, for example at (89, 100) ppm, (89, 80) ppm, (65, 100) ppm and (105, 54) ppm. The mutant exhibits stronger cellulose-pectin cross-peaks compared to the WT cell wall, as shown in various 1D cross-sections (**Fig. 5B**). For example, in the 89-ppm cellulose C4 and 65-ppm cellulose C6 cross-sections, the mutant exhibits higher cross-peaks at 100 ppm XyG C1 (xC1), Rha C1 (RC1) and GAC1 and 69 ppm (xC1/Gal C4 (GC4), GAC2/3,RC5). In the cellulose-dominant 105 ppm and 62.4 ppm cross-sections, the mutant spectrum displays higher cross-peaks to pectin at 69 ppm (x/G C4, GAC2/3, RC5), 80 ppm (A/RC2, 2_1_GAC4), 100 ppm (x/R/GAC1) and 54 ppm (COO<u>**C**</u>H_3_). Furthermore, a clear cross-peak between iC4 and 174 ppm acetyl CH_3_<u>**C**</u>OO peak appears in *gosamt1 gosamt2*, showing that there are more cellulose interactions with acetylated matrix polysaccharides (**Fig. 5A, 5B**). Given the increased 2^f^ xylan signals in the 30 ms PDSD (**Fig. 3**), it is likely this comes from xylan-cellulose interactions.

**Figure 5.**
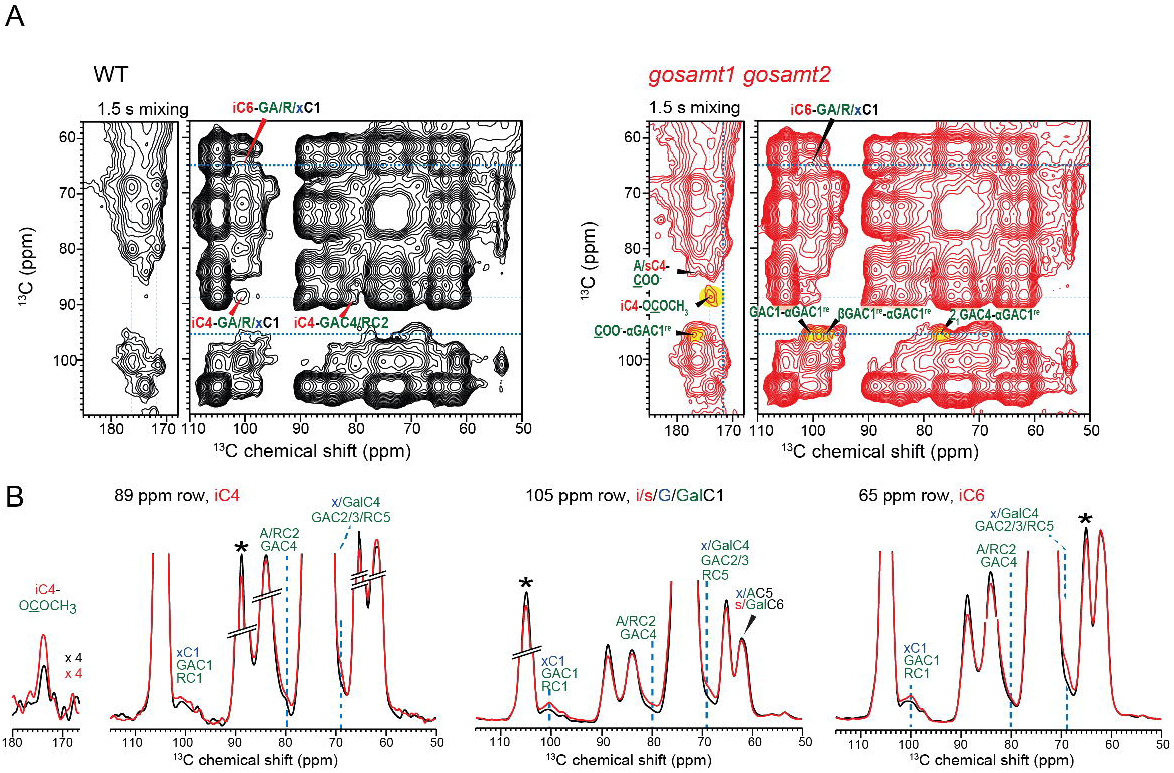
Long mixing time PDSD experiments suggest reduced methyl-esterification strengthen cellulose-pectin interactions. 1.5 s 2D ^13^C PDSD spectra of WT and *gosamt1 gosamt2* mutant cell walls. (**A**) 2D spectra of the WT and mutant cell walls, measured at 245 K under 8.8 kHz MAS using a cryoprobe. Highlighted regions in yellow show marked differences between spectra, including cross-peaks of αGAC1^re^ to 2_1_HG signals and a clear cross-peak between iC4 and 174 ppm acetyl CH_3_<u>**C**</u>OO peak. (**B**) Representative 1D ^13^C cross sections of cellulose C4 at 89 ppm and cellulose-dominant signals at 105 ppm, 65 ppm and 62 ppm. The cross sections of the two samples are scaled by the integrated area of the 55-110 ppm carbohydrate region of each 2D spectrum. Many cellulose-pectin cross-peaks (blue dashed lines) show higher intensities in the mutant than in the WT spectra, indicating that reduced esterification strengthens cellulose-pectin interactions in the wall.

Altogether these ssNMR results demonstrated the existence of 2_1_ ‘egg-box’ HG *in planta*. We identified changes in the proportion of 2_1_ and 3_1_ HG conformations, changes in polysaccharide mobility and changes in polysaccharide interactions in the mutant plants, indicating that reduced HG methylation leads to important molecular architectural changes in cell walls.

### The *gosamt1 gosamt2* mutant has clear growth and developmental phenotypes

To investigate if low HG methyl-esterification and the consequent wall changes observed at the nanoscale levels have an impact in the growth of the plants, we studied *gosamt* mutant growth phenotypes. We observed that *gosamt1* and *gosamt2* single mutants had very subtle growth penalties; however, the loss of function of *GoSAMT1* and *GoSAMT2* together leads to clear growth and developmental phenotypes. Phenotypes include dwarf plants with a cabbage-like aspect, rosettes with smaller leaves, smaller flowers, smaller siliques, and shorter inflorescence stems (**Fig. 6A**). The reduced growth of *gosamt1 gosamt2* plants is also seen in the marked differences in fresh weight and rosette diameter measurements (**Fig. 6B, 6C**). These growth phenotypes can be rescued after molecular complementation using C-terminal GFP tagged versions of GoSAMT1 and GoSAMT2 in *gosamt1 gosamt2* (**Fig. S 2A, B, C**), confirming that these phenotypes are caused by the loss of functions of *GoSAMT1* and *GoSAMT2*.

**Figure 6.**
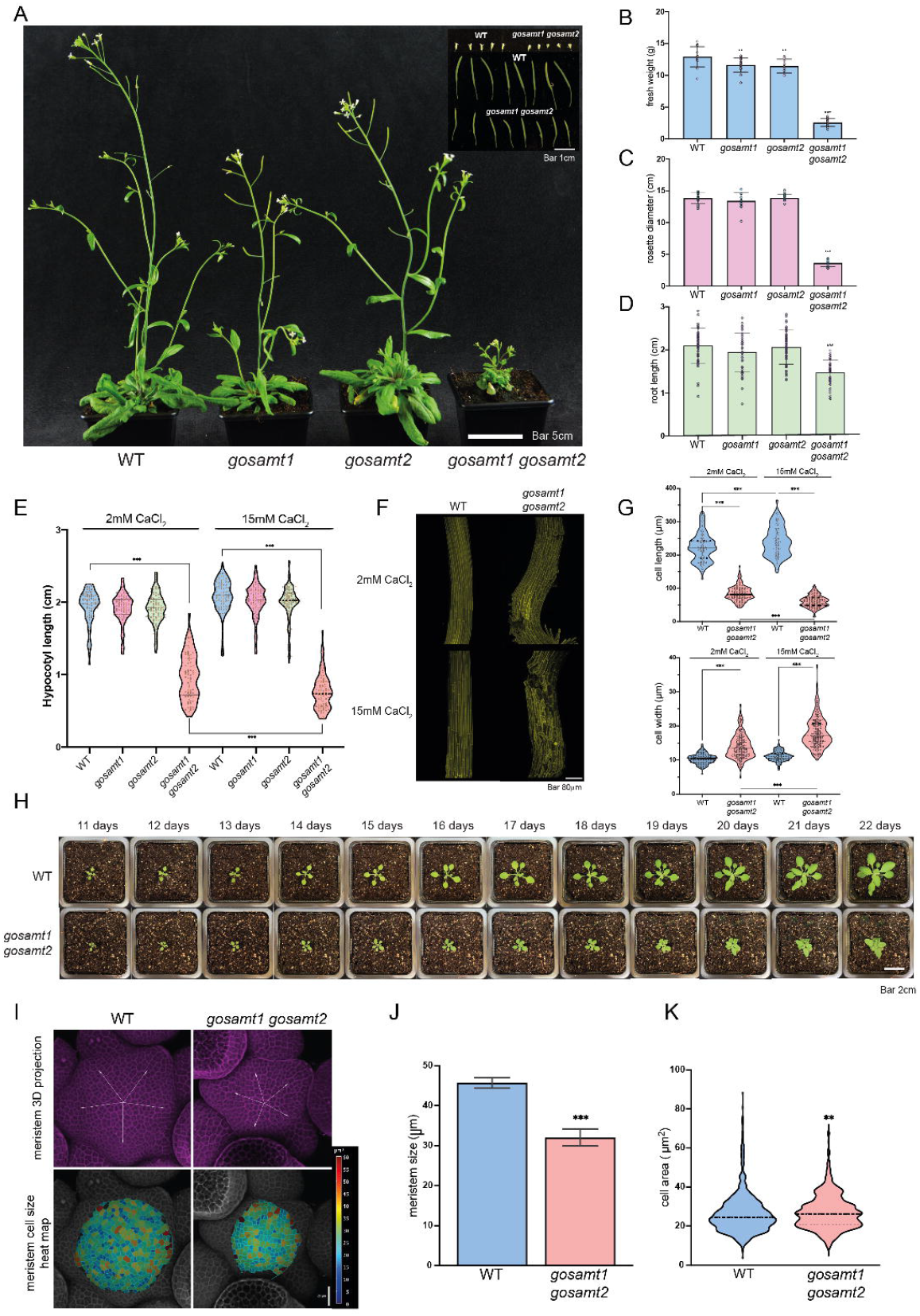
*gosamt1 gosamt2* mutants display strong growth and developmental phenotypes. (**A**) *gosamt1 gosamt2* plants display a stunted plant phenotype with shorter inflorescence stems, smaller flowers. (**B**) Fresh weight of 6-weeks-old plants. Values correspond to the mean of 15 plants per genotype. Asterisks indicate significant differences between WT and mutants defined by one-way ANOVA followed by Bonferroni’s multiple comparison test, **, P < 0.01; ***, P < 0.001. (**C**) Measurements of 6-week-old rosettes. Values correspond to the mean of 15 plants per genotype. Asterisks indicate significant differences between WT and mutants defined by one-way ANOVA followed by Bonferroni’s multiple comparison test, ***, P < 0.001. (**D**) Root length measurements of 7-day old plants. Values correspond to mean of 45 plants per genotype. Asterisks indicate significant differences between WT and mutants defined by one-way ANOVA followed by Bonferroni’s multiple comparison test, ***, P < 0.001. (**E**) Measurements of 9-day-old etiolated hypocotyls length grown in ½MS media and ½MS supplemented with CaCl_2_ to a final concentration of 15mM. Violin plots represent measurements of at least 77 plants per genotype. Asterisks indicate significant differences between mutants and the WT, and between *gosamt1 gosamt2* plants in both conditions, defined by one-way ANOVA followed by Bonferroni’s multiple comparison test, ***, P < 0.001. (**F**) Representative images of 4-days-old etiolated hypocotyls of WT and *gosamt1 gosamt2* plants expressing a plasma membrane fluorescence marker, grown in ½MS media and ½MS media with a final concentration of 15mM CaCl_2_. (**G**) Cell length and width measurements of WT and *gosamt1 gosamt2*. Violin plots represent values of at least 100 cell length and width measurements. Asterisks indicate significant differences between WT and *gosamt1 gosamt2* cell measurements defined by one-way ANOVA followed by Bonferroni’s multiple comparison test, ***, P < 0.001. (**H**) Time-course images of 11-22 day-old representative WT and *gosamt1 gosamt2* plants. (**I**) Representative images of shoot apical meristem of WT and *gosamt1* gosamt1. Meristem were stained with 0.1% propidium iodide (PI). (**J**) Measurements of meristem size in WT and *gosamt1 gosamt2* mutant, values represent the mean of N=6 for WT and N=5 for *gosamt1 gosamt2*. Asterisks indicate significant differences between WT and *gosamt1 gosamt2* cell measurements defined by unpaired t-test analysis, ***, P < 0.001. (**K**) Meristems cell area measurements, violin plot represents values of at least 548 cells. Asterisks indicate significant differences between WT and *gosamt1 gosamt2* cell measurements defined by unpaired t-test analysis, ***, P < 0.001.

We investigated to what extent *gosamt1 gosamt2* growth phenotypes reflect changes in cell expansion capacity. Time-course imaging of plants showed that leaf blades, and especially the petioles, fail to elongate at the same rate as WT plants (**Fig. 6H**). Additionally, we observed that hypocotyl and root elongation are also affected in the *gosamt1 gosamt2* mutant but not in the single mutants (**Fig. 6D, E**). The marked defects in hypocotyl elongation were observed from early after germination (**Fig. S4A**). Additionally, we investigated whether *gosamt1 gosamt2* hypocotyls presented cell adhesion phenotypes using ruthenium red dye. Ruthenium red cannot enter into WT hypocotyls, but it can enter and dye hypocotyls of mutants with cell adhesion defects ^34^. We observed *gosamt1 gosamt2* hypocotyls were stained whereas WT hypocotyls were not (**Fig. S4B**). Cell adhesion lesions were observed in the SEM (**Fig. S4C**). Remarkably, both the reduced etiolated hypocotyl elongation and poor cell adhesion phenotypes were increased when *gosamt1 gosamt2* hypocotyls were exposed to higher concentrations of Ca^2+^ (**Fig. 6E, Fig.S 4**), whereas WT plants were unaffected. These observations, together with the shorter inflorescence stems, suggest that rapid cell anisotropic growth is particularly impaired in *gosamt1 gosamt2* plants. To observe in detail hypocotyl cell anisotropic defects, we used WT and *gosamt1 gosamt2* lines expressing a fluorescent plasma membrane marker (Willis et al., 2016). We analysed the dimensions of the cells at the base of the hypocotyl, which contain the oldest and most elongated cells (Gendreau et al., 1997, Kim et al., 2015). The results show that there is a clear reduction in cell length in *gosamt1 gosamt2* when compared to WT, whereas the cell width is increased (**Fig. 6F, G**).

Shoot apical meristems are the structures which give rise to all aerial tissues in the plants, and it has been demonstrated that a fine-tuned control of pectin methylation plays a significant role in meristem maintenance and primordia formation ^9,35^. While the cell wall of the central meristem cells has been shown be highly methyl-esterified, pectin de-methyl-esterification is required for primordia formation ^36^. Therefore, since the *gosamt1 gosamt2* mutant have defects in pectin methyl-esterification we evaluated whether these plants may exhibit some changes in the shoot apical meristem. Our microscopy analysis shows that *gosamt1 gosamt2* has a smaller meristem compared to WT meristems (**Fig. 6I, J**). Interestingly, cell size is slightly increased in *gosamt1 gosamt2* mutant (**Fig. 6J**). Therefore, the number of cells is reduced in the mutant meristem, suggesting that meristematic cell division is affected.

## Discussion

SAM is the most common donor molecule for methylation reactions in all organisms. In plants, this molecule is synthesised in the cytosol, and it is utilised in many cellular pathways and locations ^37,38^. Although SAM transport into the Golgi apparatus has been proposed to be an essential step for polysaccharide methylation, this process has been difficult to study. In this work we present compelling evidence for the discovery of Golgi SAM transporters in plants. Arabidopsis mutants in these transporters suffer changes in the molecular architecture of cell walls and have strong defects in anisotropic growth of cells.

Arabidopsis possesses three *GoSAMT* genes. Our results indicate that *GoSAMT1* and *GoSAMT2*, the two most highly expressed *GoSAMTs*, play redundant roles, with the double mutant exhibiting a stronger polysaccharide methylation phenotype. However, some polysaccharide methylation remained. It will be interesting to see whether a triple mutant lacks all polysaccharide methylation, and whether this leads to lethality, though the possibility that other unrelated proteins are capable of the same function has not been ruled out.

Here, we showed that the mutation of *GoSAMTs* in Arabidopsis leads to decreased methylation of both primary cell wall pectin and secondary cell wall xylan (**Fig. 2A, B, D**), demonstrating SAM transport by a single class of transporters is important for multiple plant polysaccharide methylation processes. We know from mutants in polysaccharide methyltransferases that the absence of methylation of glucuronic acid of AGPs and of xylan does not lead to evident growth phenotypes ^3,4,39^. Any importance of methylation of GalA, GlcA, fucose and xylose in RG-II has not been previously studied ^5^, as the methyltransferases have not been identified, so it will be interesting to study RG-II methylation in *gosamt1 gosamt2* mutants in the future. On the other hand, mutants in HG pectin methylation have shown HG methylation plays an important role in cell expansion and cell adhesion ^16–18,40^. The developmental phenotypes of the *gosamt1 gosamt2* mutants are likely therefore to arise mostly from defective HG methylation. Since the process of HG methylation is controlled by a range of enzymes including numerous PMTs, PMEs and PMEIs it is difficult to study the comprehensive role of pectin methyl-esterification. The analysis of putative Golgi SAM transporter mutants is, therefore, a powerful strategy for evaluating the effects of a more global reduction in pectin methyl-esterification.

The strong decrease in pectin methyl-esterification (**Fig. 2B, 2D; Fig. 3**) led to important changes in cell wall pectin structure and conformations. Low-DM HG can form calcium-mediated HG cross-linking gel structures *in vitro*, assembling what is known as 2_1_ conformation ‘egg-box’ structures ^41^. Most of the evidence for the existence of these structures *in planta* comes from immunolabelling experiments using antibodies recognising Ca^+2^-HG complexes, and the use of genetic tools to alter the expression of PMEs and PMEIs ^36,42,43^. However, the use of non-physiological Ca^+2^ concentrations in immunolabelling experiments, and some contradictory observations showing a cell wall loosening effect instead of cell wall stiffening after PME treatments that could promote ‘egg box’ formation ^7,35,36^, raise questions about the existence and prevalence in cell walls of ‘egg box’ HG ^44^. By analysing cell walls directly by ssNMR without any pre-treatments, we identified an altered conformation of the galacturonate form of HG. This corresponds to the HG-Ca^+2^ ‘egg-box’ structure that has a two-fold screw conformation (2_1_), since the chemical shifts are similar to those reported for pectin in the different conformations *in vitro* ^32^. Interestingly, reduced HG methylation in *gosamt1 gosamt2* led to an increase in the proportion of the of 2_1_ HG-Ca^+2^ ‘egg-box’ vs three-fold (3_1_) HG conformations (**Fig. 3**), and an increase in HG reducing ends. The *gosamt1 gosamt2* mutant therefore contains more abundant and short ‘egg-box’ structures (**Fig. 5**). Polysaccharide mobility changes were observed using DIPSHIFT NMR, indicating the HG backbone is more rigid in the mutant, whereas RG-I side chains appear to be more mobile (**Fig. 4**). Together, these results are compelling evidence of the existence of ‘egg-box’ structures that crosslink HG backbones in plant cell walls. Moreover, the formation of ‘egg-box’ HG is promoted by low methyl-esterification.

Important changes in the cell wall polymer interactions were observed in our ssNMR experiments. We detected stronger signals of the cellulose-bound form of two-fold screw xylan in *gosamt1 gosamt2* (**Fig. 3**). It is not expected that reduced pectin methyl-esterification leads to higher xylan content, and there are no significant changes in the content of xylose in our sugar composition analysis (**Fig. 2C**), or changes in xylan signals in our quantitative 1D NMR experiments (**Fig. 2D**). On the other hand, changes in pectin mobility could influence the capacity of xylan to interact with cellulose because pectin – xylan interactions may occur in cell walls ^45–47^. An increase of pectin in a gelled state could facilitate more xylan binding to cellulose, increasing the NMR signal of two-fold screw xylan in *gosamt1 gosamt2*. This idea is supported by the increased signal of the cross-peak between cellulose iC4 89 ppm and acetyl CH_3_<u>**C**</u>OO^−^ 174 ppm in our 1.5s PDSDs spectra, which is likely to arise from acetylated xylan due to its strong capacity to bind to cellulose ^48^. The increased pectin-cellulose cross-peaks (**Fig. 5B**), suggest that changes in pectin properties also increased the contacts between these two polysaccharides.

The cell wall changes detected at the nanoscale level in the *gosamt1 gosamt2* mutant help evaluate the complex relationship between pectin methyl-esterification and cell expansion. Higher levels of pectin methyl-esterification have been correlated with rapid anisotropic growth in inflorescence stems ^49^ and dark-grown hypocotyls ^50^. In the *gosamt1 gosamt2* mutants which have reduced pectin methyl-esterification, reduced anisotropic growth is reflected in the shortened inflorescence stems, roots, petioles, and dark-grown hypocotyls (**Fig. 6**). Moreover, an increased concentration of calcium in the media further reduced the capacity of hypocotyl cells to expand and increased the severity of cell adhesion phenotypes. Perhaps higher calcium could interact with the unusually low esterification pectin in the mutant, leading to increased calcium ‘egg-box’ formation, increasing pectin rigidity, or altering pectin interaction with cellulose fibrils. Cellulose anisotropy has also been proposed as a major factor dictating how cells expand ^51,52^. Even though we did not observe changes in cellulose structure at the nano-scale level, it would be interesting to evaluate changes in cellulose arrangement at the meso-scale level, because pectin methylation mutant *qua2* shows important changes in cellulose deposition ^16^. Changes in the HG content were observed in *gosamt1 gosamt2* (**Fig. 2C, D; Fig. 3**). This has also been observed in PMT mutants ^16,17,20^. Proper pectin methyl-esterification may be required for appropriate pectin synthesis, for example to prevent gelling in the Golgi, or to avoid electrostatic repulsion due to abundant GalA negative charges. Additionally, low-DM pectin could serve as a substrate for PGAses *in muro* ^8,11^. However, the observed decrease in HG quantity in the *gosamt1 gosamt2* mutants seems unlikely solely to underlie the growth and developmental phenotypes, since greater HG reductions in glucuronate 4-epimerase mutants ^53^ do not result in strong growth and developmental phenotypes.

Pectin methyl-esterification has been proposed to be a key factor for meristem maintenance and primordia formation ^35,36^. Here we showed that *gosamt1 gosamt2* mutant has a significant decrease in the size of the meristem by having fewer cells, supporting the notion that appropriate pectin methyl-esterification is required for meristem cell division. Similar meristem observations were made in a mutant in an orthologous MFS_5 in rice ^54^. These meristem changes will contribute to the growth and developmental phenotypes of the *gosamt1 gosamt2* mutant.

The most obvious changes in *gosamt1 gosamt2* cell walls likely arise directly from differences in the degree of methyl-esterification of HG and the ensuing changes in its mechanical properties, yet we need to consider additional possibilities that could influence the mutant plant phenotypes. For example, wall stress signals may be triggered in *gosamt1 gosamt2*. Recently, it was reported that stress response genes were upregulated in the PMT *qua2* mutants ^16^. Oligogalacturonides have been demonstrated to act as potent DAMPs ^55,56^, and reduced pectin methyl-esterification promotes PGAse activity, suggesting that higher accumulation of OGAs could occur in *gosamt1 gosamt2*. Indeed, the ssNMR experiments suggested there are increases in HG reducing ends in the mutant. Alternatively, there could be altered activation of the cell wall receptor WAK1 because it binds more strongly to ‘egg-box’ structures ^57^. Lastly, other effects related to SAM metabolism may need to be considered. Reduced incorporation into the Golgi apparatus could lead to changes in the cytosolic SAM pool. It has been reported that an uncontrolled balance of SAM/SAH leads to changes in DNA methylation ^58^, however, this balance is tightly regulated by SAHH enzymes ^59^.

This work allowed us to establish the important role of these putative transporters in the process of polysaccharide methylation. While future experiments assaying *in vitro* activity are required to understand the transport mechanisms of GoSAMTs, the discovery of these transporters is a significant step in understanding Golgi synthesised polysaccharide methylation.

## Materials and methods

### Plant material

The T-DNA insertion lines analysed in this study were in the Arabidopsis (*Arabidopsis thaliana*) Col-0 background. The *gosamt1* (SALK_054431) and *gosamt2* (GABI_480G06) were provided by the Nottingham Arabidopsis Stock Centre. Homozygous mutants were identified by PCR genotyping (for oligonucleotide sequences, see Supplemental Table 1). To identify knock-out mutants, RNA was extracted from homozygous mutant leaves using the RNAeasy Mini Kit (Qiagen). The extracted RNA was treated with DNase (RQ1 RNase-Free DNase, Promega). cDNA was generated using reverse transcriptase (SuperScript II Reverse Transcriptase, Invitrogen), and RT-qPCR was performed using oligonucleotides listed in Supplemental Table 1. Plants were grown on soil (Advance M2, ICL Levington) in a cabinet maintained at 21 °C, with a 16 h light, 8 h dark photoperiod. For in vitro growth seeds were surface sterilized and sown on solidified basal ½MS medium (2.2 g/L; M5519, Sigma Aldrich) containing 1.0% (w/v) sucrose and 0.1% (w/v) MES, and the pH was adjusted to 5.8 using KOH and HCl. The sown seeds were stratified for 2 d at 4°C and incubated at 21°C under white light (150 μmol m^−2^ s^−1^) with a 16-h-light/8-h-dark cycle.

### Molecular cloning and plant transformation

Constructs were obtained using Golden Gate DNA assembly protocol ^60^. Sequences were optimized for the removal of the enzyme restriction sites *Bsa*I, *Bpi*I, *Esp*3I, and *Dra*III. The stop codons were also removed from CDSs to allow GFP C-terminal fusions. The restriction enzyme *Bsa*I was used for level 1 assembly and *Bpi*I was used for level 2 assembly. 1752 bp upstream the start codon of *At1g64650* were used as *_pro_GoSAMT1*, 796 bp intergenic region of *At4g27720* and *At4g27710* was as *_pro_GoSAMT2*. For molecular complementation experiments, level 1 assemblies were made using GoSAMTs promoters and CDSs were fused to eGFP followed by a NOS terminator. Level 2 assemblies were made using FAST selection marker ^61^ in position one, *_pro_GoSAMT-GoSAMT-GFP-NOST* in position 2 and the Golgi marker GmMan1-mCherry expressed under CaMV 35S promoter ^62^ in position 3. Arabidopsis transformation was performed using floral dip as previously described ^63^, and transformants were selected by fluorescent screening.

### Monosaccharide composition analysis

0.5mg AIR samples were hydrolysed in 2 M trifluoroacetic acid (TFA) for 1h at 120°C. Following evaporation under vacuum for the removal of TFA, the samples were resuspended in 400 μL water. The resulting monosaccharides were detected, and the sugar composition was determined by HPAEC-PAD using an ICS-3000+ fitted with a pulsed amperometric detector (Thermo Fisher Scientific). Monosaccharides included in the hydrolysate were separated on a Dionex CarboPac™ PA-20 column (3×150mm; Thermo Fisher Scientific).

### Subcellular localisation

14 days-old leaves of stable transgenic Arabidopsis lines expressing *_pro_GoSAMT1:GoSAMT1-eGFP* or *_pro_GoSAMT:GoSAMT2-eGFP* and Golgi marker GmMan1-mCherry were scanned using a Leica SP8 confocal microscope with HC PL APO CS2 63x/1.20 water lens. eGFP and mCherry signals were detected in sequential mode. Laser line 488 and hybrid detector (HyD) was used for eGFP detection, and laser line 552 and photomultiplier tube (PMT) detector was used for mCherry detection.

### Etiolated hypocotyl and meristem measurements

For etiolated hypocotyl growth, seeds were plated on ½MS media or ½MS media supplemented with 12.5mM CaCl_2_. Seeds were stratified for 2 d at 4°C and plates were kept in light until the radicle breaks the endosperm and then covered with two layers of aluminium foil for 3-9 days. Hypocotyls were scanned using EPSON Perfection V600 Photo and length were analysed using FIJI ^64^. For hypocotyl cell analysis, a plasma membrane fluorescence marker myr-YFP (an acylated YFP) ^65^ was introduced into WT and *gosamt1 gosamt2* backgrounds via agrobacterium mediated transformation. Etiolated hypocotyls of the transgenic lines were grown in MS plates or MS plates supplemented with 12.5mM for 4 days. The cells at the bottom part of the hypocotyl (close to the root) were scanned in Zeiss LSM700 with a 10 × dry objective. 3D images were constructed in MorphoGraphX. The cell length and width were measured in Fiji. Cryo-scanning electron microscopy (cryo-SEM) was carried out on a Zeiss EVO HD15 SEM fitted with a Quorum PT3010T cryo system (Quorum Technologies, Lewes, UK). The cryo preparation and imaging procedure was carried out as described by Wightman et al. ^66^ with the following modifications: A 12 mm diameter adhesive carbon tab was fixed to the brass stub using a 3:1 mixture of Tissue-Tec (Scigen Scientific, USA) and colloidal graphite in order to prevent the tabs falling off the stub during the plunge freezing. 4 DAG dark grown hypocotyls of WT and *gosamt1 gosamt2* grown in ½MS media or ½MS media supplemented with 12.5mM CaCl_2_ were placed on the adhesive tab and then plunge frozen in slush nitrogen and then transferred under vacuum to the cryo prep chamber. The frozen samples were then sputter coated to a thickness of 5 nm using a Gold/Palladium target and then transferred to the cryo-stage in the SEM chamber. Images were taken on the SEM using a gun voltage of 25 kV, I probe size of 38 pA, a SE detector and a working distance of approx 13 mm. Morphological analysis of the shoot apical meristem (SAM) was performed on plants shortly after bolting. The shoot apices of WT and *gosamt1/2* were cut, and the mature flowers and flower buds were removed. The dissected meristems were stained with 0.1% propidium iodide (PI) for 5 min in a square box containing fresh MS medium (Duchefa Biochemie-MS basal salt mixture) supplemented with vitamins and 1% sucrose. After briefly rinsing in water, the SAMs were scanned in Zeiss LSM700 with 20 × NA 1.0 water dipping objective. Laser excitation was 488 nm for PI. The SAM size was analysed in Fiji by measuring the radius of the 3D projections of the confocal z-stacks. For cell size analysis, the confocal z-stacks were segmented in MorphoGraphX ^67^, and the size of the L1 layer cells were quantified.

### Xylan digestion and DASH analysis

250μg of AIR from basal stems was incubated overnight in 0.1 M ammonium acetate buffer (pH 5.5) with an excess of Neocallimastix patriciarum Xyn11A xylanase ^68^ at 21°C. The derivatisation of oligosaccharides with 8-aminopyrene-1,4,6-trisulfonate (APTS) was performed according to previously developed protocols ^29^. A set of 7 fuorophore (DY-481XL-NHS ester)-labelled amino acids and peptides was used as electrophoretic mobility standards (Asp–Asp–Asp–Asp; Asp–Asp–Asp; Glu–Glu; Cysteic acid; l-2-Aminoadipic acid; Glycine; Gly–Gly–Gly) to align the electropherograms. These electrophoretic mobility standards were mixed with each sample prior to DASH separation serving as internal mobility markers. DASH data generated by the DNA sequencer were processed with the DASHboard software ^29^.

### Methanol release assay of AIR samples

1mg of AIR samples were treated with 100 μL 0.2M NaOH for an hour at 4 degrees to release methanol from methyl-esterified pectin, and then neutralised with 100 μL 0.2M HCL and buffered with 200 μL of 50mM Tris-HCL pH7.0. Methanol released was measured using MBTH method described in^69^. 100μL of 100mM Tris-HCL pH7.0, 400μL (3 mg/mL) of methylbenzothiazolinone-2-hydrazone (MBTH, Sigma M8006−1G), 200μL of sample and 0.5μI of alcohol oxidase (E.C. 1.1.3.13, Sigma, A2404) were mixed to oxidize the methanol. After addition of the alcohol oxidase, samples were incubated for 20 min at 30°C. 200 μL of a solution containing 5 mg/mL each of dodecahydrated ferric ammonium sulfate and sulfamic acid was added and the solution was incubated for 20 min at room temperature. 600 μL of water was added the tubes were vortexed. Absorbance at 620 nm was measured to plot a standard curve of concentration and absorbance.

### Growth of ^13^C-labeled Arabidopsis primary cell walls

WT and *gosamt1 gosamt2* plants were grown in a custom-designed growth chamber and were enriched in ^13^C using CO_2_ as previously described ^70^. Specifically, hydroponics solution (2 mM MgSO_4_, 2 mM calcium], 50 mM FeEDTA, 5 mMKNO_3_, 2.5 mM K_2_HPO4/KH_2_PO4 pH 5.5, 70 mM H_3_BO_4_, 14 mM MnCl_2_, 0.5 mM CuSO_4_, 1 mM ZnSO_4_, 0.2 mM NaMoO_4_, 10 mM NaCl and 0.1 mM CoCl_2_) was poured into the chamber twice a week, and the excess media was removed after pouring the media. *Arabidopsis* seedlings were placed on rockwool that was covered with a foil pierced with holes and placed in the growth chamber. Compressed air was scrubbed of CO_2_ using calcium oxide, then ^13^C-enriched CO_2_ was mixed in at a concentration of 500 p.p.m. before entering the growth chamber. Plants were grown for 6 weeks at 22°C and 60-70% humidity in cycles of 16 h light and 8 h dark.

### Solid-state NMR spectroscopy

All solid-state NMR spectra were measured on 800 MHz (18.8 T), 700 MHz (16.4 T) and 600 MHz (14.1 Tesla) Bruker Advance II spectrometers using 3.2 mm MAS probes. Typical radiofrequency (rf) field strengths were 40-62.5 kHz for ^13^C and 50-83 kHz for ^1^H. Two-pulse phase-modulated (TPPM) ^1^H decoupling was applied during acquisition. All ^13^C chemical shifts were externally referenced to the adamantane CH_2_ peak at 38.48 ppm on the tetramethylsilane (TMS) scale.

The 1D DP spectra of ^13^C labelled fresh plants shown in **Figure 2** were taken at 290 K, 10.1 kHz MAS and had 1024 acquisitions each with a 20 s recycle delay.

Two sets of 2D proton-driven ^13^C-^13^C spin diffusion (PDSD) spectra were measured. One set of spectra were measured at 275 K using a short mixing time on an Efree probe, whereas the second set of spectra were measured at 245 K using a long mixing time on a Biosolids Cryoprobe. The higher-temperature spectra were measured at 800 MHz under 11 kHz MAS, using a 100 ms mixing time, a recycle delay of 1.8 s and a CP contact time of 500 μs. This experiment mainly detects intramolecular cross-peaks, especially for the 50-60% ^13^C-labeled sample. The spectral widths were 249 ppm (50 kHz) and 124 ppm (25 kHz) for the direct and indirect dimensions, respectively. The indirect dimension has 120 t_1_ increments and a maximum evolution time of 4.8 ms, and 256 scans and 288 scans were acquired for the WT and mutant samples, respectively. The lower-temperature 2D PDSD spectra were measured on the 70-80% ^13^C-labeled samples at 600 MHz under 8.8 kHz MAS, using a mixing time of 1.5 s, a recycle delay of 1.5 s, and a CP contact time of 500 μs. The cryoprobe enhanced the spectral sensitivity 3-fold compared to the regular Efree probe. The 1.5 s mixing PDSD experiments detect intermolecular cross-peaks under conditions where the matrix polysaccharides are significantly immobilized. The spectral widths were 394 ppm (60 kHz) and 181 ppm (27 kHz) for the direct and indirect dimensions, respectively. The indirect dimension has 273 t_1_ increments and a maximum evolution time of 10 ms, and 192 scans and 160 scans were acquired for the WT and mutant samples.

Several types of 2D ^13^C-^1^H dipolar chemical-shift (DIPSHIFT) correlation experiments ^71,72^ were carried out on the 800 MHz spectrometer to investigate the polysaccharide mobility in the WT and mutant cell walls. These experiments differ in whether the initial ^13^C magnetization was created by CP or DP and whether a short or long recycle delay was used. The quantitative DIPSHIFT experiment was conducted using ^13^C DP and a long recycle delay of 20 s to obtain quantitative intensities that reflect the motional amplitudes of all polysaccharides. These spectra were measured under 7.8 kHz MAS at 275 K. The CP-DIPSHIFT experiment preferentially detects the mobility of more rigid polysaccharides, and used a 2 s recycle delay and a 500-μs CP contact time. The spectra were measured under 7.8 kHz MAS at 293 K. A DP-DIPSHIFT experiment with an intermediate recycle delay of 12 s was also conducted to measure the dynamics of most matrix polysaccharides while suppressing the signals of the rigid cellulose. For all DIPSHIFT experiments, the dipolar coupling was measured in a doubled fashion by keeping the ^1^H homonuclear decoupling period constant at one rotor period while shifting the ^13^C 180° pulse during the rotor period ^72^. The FSLG pulse sequence ^73^ with a ^1^H transverse field strength of 73-75 kHz was used for ^1^H homonuclear decoupling. The theoretical FSLG scaling factor of 0.577 was verified using the model peptide formyl-Met-Leu-Phe-OH ^74^. The measured dipolar couplings were divided by the scaled rigid-limit value of 26.2 kHz to obtain the C-H bond order parameter, S_CH_. All experiments were measured with dipolar-doubled version.

## Author contribution

H.T., A.O., M.H. and P.D. conceived and designed the study. H.T. conducted most of the molecular genetic, microscopy and biochemical experiments, assisted by W.Y., J.J.L., A.E.P., I.Y., J.P.P-R., O.M.T. and S.S-A. ssNMR experiments were conducted mostly by P.P. with contribution of R.D. Data analysis and interpretation was conducted by H.T., P.P., A.O., M.H. and P.D. The paper was written by H.T. and P.D. with contributions from all authors.

## Acknowledgments

The characterisation of *gosamt* mutants was supported as part of The Center for Lignocellulose Structure and Formation, an Energy Frontier Research Center funded by the U.S. Department of Energy (DOE), Office of Science, Basic Energy Sciences (BES), under Award # DE-SC0001090. This study made use of NMR spectrometers at the MIT-Harvard Center for Magnetic Resonance, which is supported by NIH grant P41 GM132079. Initial gene identification, mutant isolation and preliminary pectin methylation studies were done by H.T. and P.D. under grant EPSRC/BBSRC OpenPlant (BB/L014130/1) and A.O., J.P.P-R and S.S-A supported by Fondo de Areas Prioritarias- Centro de Regulacion del Genoma-15090007, FONDECYT 1190695 and FONDECYT 1201467.

We thank Dr. Federico López-Hernández for his advice on dark-grown hypocotyl experiments, Dr. Raymond Wightman for his helpful support with microscopy experiments, Dr. Louis Wilson and Dr. Nadine Anders for their helpful suggestions during manuscript writing.

## Supplemental Figure Legends

**Supplemental Figure 1. Arabidopsis GoSAMTs show similarity to CeSAMT1 and they are present throughout the plant kingdom. (A)** Sequence identity and similarity matrix generated based on MUSCLE alignments of whole length protein sequences of CeSAMT1 and Arabidopsis GoSAMTs. **(B)** CeSAMT1 and Arabidopsis GoSAMTs topology using TOPCONS ^75^. **(C)** Phylogenetic tree of MFS_5 proteins from *Arabidopsis thaliana* (At), *Amborella thricopoda* (Atr), *Caenorhabditis elegans, Chlamydomona reinhardtii, Klebsormidium nitens, Homo* sapiens, *Micromonas commoda, Oryza sativa* (Os), *Ostreococcus lucimarinus*, Picea *abies* (PAB), *Physcomitrella patens* (Pp) *Selaginella moellendorffii* (SMO), *Solanum lycopersicum* (Solyc). Arabidopsis sequences are highlighted in red. Phylogenetic tree was generated using Molecular Evolutionary Genetics Analysis MEGA X and visualized by FigTree 1.4.2.

**Supplemental Figure 2. Complementation of *gosamt1 gosamt2* mutant. (A)** Pictures of adult plants of WT, *gosamt* mutants and *gosamt1 gosamt2* mutant molecular complemented lines. **(B)** Western blot, expression analysis of GoSAMT1-GFP and GoSAMT2-GFP complemented lines using anti-GFP. **(C)** Box and whiskers plot representing plant fresh weight of the different genotypes. **(D)** Ratio of released Xyl_4_GlcA and Xyl_4_^Me^GlcA products after endoxylanase treatment of basal stem AIR, coupled to capillary electrophoresis experiments of secondary cell wall xylan of WT, *gosamt1*, *gosamt2*, *gosamt1 gosamt2* and molecular complemented lines. Values correspond to the mean of 3 biological replicates. **(D)** Methanol release experiments of leaf AIR of WT, *gosamt1*, *gosamt2*, *gosamt1 gosamt2* and molecular complemented lines. Values correspond to the mean of 3 biological replicates.

**Supplemental Figure 3. CP-DIPSHIFT experiment.** CP-DIPSHIFT curves of WT (black) and *gosamt1 gosamt2* (red) cell walls measured at 293 K. Best-fit ^13^C-^1^H dipolar coupling values (scaled by FSLG) and S_CH_ order parameters are given in each panel. The experiment was conducted with C-H dipolar doubling, a CP contact time of 500 μs under 7.8 kHz MAS. The 101 ppm, 99.0 ppm, and 79.8 ppm peaks of pectin backbones are more rigid in the mutant than in the WT cell wall.

**Supplemental Figure 4. *gosamt1 gosamt2* display strong cell elongation and cell adhesion phenotypes in etiolated hypocotyls. (A)** Quantification of etiolated hypocotyls of WT, *gosamt1*, *gosamt2* and *gosamt1 gosamt2*, one to five days after germination (DAG) grown etiolated hypocotyls length grown in ½MS media and ½MS supplemented with CaCl_2_ to a final concentration of 15mM. n ≥73 seedlings per genotype per time point. **(B)** WT and *gosamt1 gosamt2* hypocotyls etiolated hypocotyls length grown in ½MS media and ½MS supplemented with CaCl_2_ to a final concentration of 15mM stained with ruthenium red. Bar 5mm **(C)** Scanning electron microscopy (SEM) of WT and in MS and in MS supplemented with CaCl_2_ to a final concentration of 15 mM. Bar 100μm.

## References

1 Temple, H., Saez-Aguayo, S., Reyes, F. C. & Orellana, A. The inside and outside: topological issues in plant cell wall biosynthesis and the roles of nucleotide sugar transporters. Glycobiology 26, 913–925, doi:10.1093/glycob/cww054 (2016).

2 Reyes, F. & Orellana, A. Golgi transporters: opening the gate to cell wall polysaccharide biosynthesis. Current opinion in plant biology 11, 244–251 (2008).

3 Urbanowicz, B. R. et al. 4-O-methylation of glucuronic acid in Arabidopsis glucuronoxylan is catalyzed by a domain of unknown function family 579 protein. Proc Natl Acad Sci U S A 109, 14253–14258, doi:10.1073/pnas.1208097109 (2012).

4 Temple, H. et al. Two members of the DUF579 family are responsible for arabinogalactan methylation in Arabidopsis. Plant Direct 3, e00117, doi:10.1002/pld3.117 (2019).

5 O’Neill, M. A. et al. Locating Methyl-Etherified and Methyl-Esterified Uronic Acids in the Plant Cell Wall Pectic Polysaccharide Rhamnogalacturonan II. SLAS Technol 25, 329–344, doi:10.1177/2472630320923321 (2020).

6 Atmodjo, M. A., Hao, Z. & Mohnen, D. Evolving views of pectin biosynthesis. Annu Rev Plant Biol 64, 747–779, doi:10.1146/annurev-arplant-042811-105534 (2013).

7 Levesque-Tremblay, G., Pelloux, J., Braybrook, S. A. & Müller, K. Tuning of pectin methylesterification: consequences for cell wall biomechanics and development. Planta 242, 791–811 (2015).

8 Daher, F. B. & Braybrook, S. A. How to let go: pectin and plant cell adhesion. Front Plant Sci 6, 523, doi:10.3389/fpls.2015.00523 (2015).

9 Palin, R. & Geitmann, A. The role of pectin in plant morphogenesis. Biosystems 109, 397–402, doi:10.1016/j.biosystems.2012.04.006 (2012).

10 Goldberg, R., Morvan, C., Jauneau, A. & Jarvis, M. in Progress in biotechnology Vol. 14 151–172 (Elsevier, 1996).

11 Xiao, C., Somerville, C. & Anderson, C. T. POLYGALACTURONASE INVOLVED IN EXPANSION1 functions in cell elongation and flower development in Arabidopsis. Plant Cell 26, 1018–1035, doi:10.1105/tpc.114.123968 (2014).

12 Phyo, P., Wang, T., Xiao, C., Anderson, C. T. & Hong, M. Effects of Pectin Molecular Weight Changes on the Structure, Dynamics, and Polysaccharide Interactions of Primary Cell Walls of Arabidopsis thaliana: Insights from Solid-State NMR. Biomacromolecules 18, 2937–2950, doi:10.1021/acs.biomac.7b00888 (2017).

13 Yang, Y., Anderson, C. T. & Cao, J. POLYGALACTURONASE45 cleaves pectin and links cell proliferation and morphogenesis to leaf curvature in Arabidopsis thaliana. The Plant Journal (2021).

14 Zhang, G. F. & Staehelin, L. A. Functional compartmentation of the Golgi apparatus of plant cells: immunocytochemical analysis of high-pressure frozen-and freeze-substituted sycamore maple suspension culture cells. Plant physiology 99, 1070–1083 (1992).

15 Goubet, F. & Mohnen, D. Subcellular localization and topology of homogalacturonan methyltransferase in suspension-cultured Nicotiana tabacum cells. Planta 209, 112–117, doi:10.1007/s004250050612 (1999).

16 Du, J. et al. Mutations in the Pectin Methyltransferase QUASIMODO2 Influence Cellulose Biosynthesis and Wall Integrity in Arabidopsis. Plant Cell 32, 3576–3597, doi:10.1105/tpc.20.00252 (2020).

17 Mouille, G. et al. Homogalacturonan synthesis in Arabidopsis thaliana requires a Golgi-localized protein with a putative methyltransferase domain. Plant J 50, 605–614, doi:10.1111/j.1365-313X.2007.03086.x (2007).

18 Krupkova, E., Immerzeel, P., Pauly, M. & Schmulling, T. The TUMOROUS SHOOT DEVELOPMENT2 gene of Arabidopsis encoding a putative methyltransferase is required for cell adhesion and co-ordinated plant development. Plant J 50, 735–750, doi:10.1111/j.1365-313X.2007.03123.x (2007).

19 Miao, Y., Li, H. Y., Shen, J., Wang, J. & Jiang, L. QUASIMODO 3 (QUA3) is a putative homogalacturonan methyltransferase regulating cell wall biosynthesis in Arabidopsis suspension-cultured cells. J Exp Bot 62, 5063–5078, doi:10.1093/jxb/err211 (2011).

20 Kim, S. J., Held, M. A., Zemelis, S., Wilkerson, C. & Brandizzi, F. CGR2 and CGR3 have critical overlapping roles in pectin methylesterification and plant growth in Arabidopsis thaliana. Plant J 82, 208–220, doi:10.1111/tpj.12802 (2015).

21 Ibar, C. & Orellana, A. The import of S-adenosylmethionine into the Golgi apparatus is required for the methylation of homogalacturonan. Plant Physiol 145, 504–512, doi:10.1104/pp.107.104679 (2007).

22 Nikolovski, N. et al. Putative glycosyltransferases and other plant Golgi apparatus proteins are revealed by LOPIT proteomics. Plant Physiol 160, 1037–1051, doi:10.1104/pp.112.204263 (2012).

23 Nikolovski, N., Shliaha, P. V., Gatto, L., Dupree, P. & Lilley, K. S. Label-free protein quantification for plant Golgi protein localization and abundance. Plant Physiol 166, 1033–1043, doi:10.1104/pp.114.245589 (2014).

24 Yan, N. Structural biology of the major facilitator superfamily transporters. Annual review of biophysics 44, 257–283 (2015).

25 Tejada-Jimenez, M., Galvan, A. & Fernandez, E. Algae and humans share a molybdate transporter. Proc Natl Acad Sci U S A 108, 6420–6425, doi:10.1073/pnas.1100700108 (2011).

26 Wohlschlager, T. et al. Methylated glycans as conserved targets of animal and fungal innate defense. Proc Natl Acad Sci U S A 111, E2787–2796, doi:10.1073/pnas.1401176111 (2014).

27 Obayashi, T., Aoki, Y., Tadaka, S., Kagaya, Y. & Kinoshita, K. ATTED-II in 2018: a plant coexpression database based on investigation of the statistical property of the mutual rank index. Plant and Cell Physiology 59, e3–e3 (2018).

28 Schmid, M. et al. A gene expression map of Arabidopsis thaliana development. Nature genetics 37, 501–506 (2005).

29 Li, X. et al. Development and application of a high throughput carbohydrate profiling technique for analyzing plant cell wall polysaccharides and carbohydrate active enzymes. Biotechnol Biofuels 6, 94, doi:10.1186/1754-6834-6-94 (2013).

30 Wang, T., Park, Y. B., Cosgrove, D. J. & Hong, M. Cellulose-pectin spatial contacts are inherent to never-dried *Arabidopsis thaliana* primary cell walls: evidence from solid-state NMR. Plant Physiol. 168, 871–883 (2015).

31 Hediger, S., Emsley, L. & Fischer, M. Solid-state NMR characterization of hydration effects on polymer mobility in onion cell-wall material. Carbohydr. Res. 322, 102–112, doi:Doi 10.1016/S0008-6215(99)00195-0 (1999).

32 Jarvis, M. C. & Apperley, D. C. Chain Conformation in Concentrated Pectic Gels - Evidence from C-13 Nmr. Carbohyd Res 275, 131–145, doi:Doi 10.1016/0008-6215(95)00033-P (1995).

33 Wang, T., Phyo, P. & Hong, M. Multidimensional solid-state NMR spectroscopy of plant cell walls. Solid State Nucl. Magn. Reson. 78, 56–63, doi:10.1016/j.ssnmr.2016.08.001 (2016).

34 Kohorn, B. D. et al. Pectin Dependent Cell Adhesion Restored by a Mutant Microtubule Organizing Membrane Protein. Plants (Basel) 10, doi:10.3390/plants10040690 (2021).

35 Peaucelle, A. et al. Arabidopsis phyllotaxis is controlled by the methyl-esterification status of cell-wall pectins. Curr Biol 18, 1943–1948, doi:10.1016/j.cub.2008.10.065 (2008).

36 Peaucelle, A. et al. Pectin-induced changes in cell wall mechanics underlie organ initiation in Arabidopsis. Curr Biol 21, 1720–1726, doi:10.1016/j.cub.2011.08.057 (2011).

37 Palmieri, L. et al. Molecular identification of an Arabidopsis S-adenosylmethionine transporter. Analysis of organ distribution, bacterial expression, reconstitution into liposomes, and functional characterization. Plant physiology 142, 855–865 (2006).

38 Bouvier, F. et al. Arabidopsis SAMT1 defines a plastid transporter regulating plastid biogenesis and plant development. The Plant Cell 18, 3088–3105 (2006).

39 Lee, C. et al. Three Arabidopsis DUF579 domain-containing GXM proteins are methyltransferases catalyzing 4-O-methylation of glucuronic acid on xylan. Plant and Cell Physiology 53, 1934–1949 (2012).

40 Bouton, S. et al. QUASIMODO1 encodes a putative membrane-bound glycosyltransferase required for normal pectin synthesis and cell adhesion in Arabidopsis. Plant Cell 14, 2577–2590, doi:10.1105/tpc.004259 (2002).

41 Cao, L., Lu, W., Mata, A., Nishinari, K. & Fang, Y. Egg-box model-based gelation of alginate and pectin: A review. Carbohydrate Polymers, 116389 (2020).

42 Liners, F., Thibault, J.-F. & Van Cutsem, P. Influence of the degree of polymerization of oligogalacturonates and of esterification pattern of pectin on their recognition by monoclonal antibodies. Plant physiology 99, 1099–1104 (1992).

43 Draye, M. & Van Cutsem, P. Pectin methylesterases induce an abrupt increase of acidic pectin during strawberry fruit ripening. Journal of plant physiology 165, 1152–1160 (2008).

44 Hocq, L., Pelloux, J. & Lefebvre, V. Connecting homogalacturonan-type pectin remodeling to acid growth. Trends in Plant Science 22, 20–29 (2017).

45 Ralet, M.-C. et al. Xylans provide the structural driving force for mucilage adhesion to the Arabidopsis seed coat. Plant Physiology 171, 165–178 (2016).

46 Broxterman, S. E. & Schols, H. A. Characterisation of pectin-xylan complexes in tomato primary plant cell walls. Carbohydrate polymers 197, 269–276 (2018).

47 Biswal, A. K. et al. Working towards recalcitrance mechanisms: increased xylan and homogalacturonan production by overexpression of GAlactUronosylTransferase12 (GAUT12) causes increased recalcitrance and decreased growth in Populus. Biotechnology for biofuels 11, 1–26 (2018).

48 Simmons, T. J. et al. Folding of xylan onto cellulose fibrils in plant cell walls revealed by solid-state NMR. Nat Commun 7, 13902, doi:10.1038/ncomms13902 (2016).

49 Phyo, P. et al. Gradients in Wall Mechanics and Polysaccharides along Growing Inflorescence Stems. Plant Physiol 175, 1593–1607, doi:10.1104/pp.17.01270 (2017).

50 Daher, F. B. et al. Anisotropic growth is achieved through the additive mechanical effect of material anisotropy and elastic asymmetry. Elife 7, e38161 (2018).

51 Zhang, Y. et al. Molecular insights into the complex mechanics of plant epidermal cell walls. Science 372, 706–711, doi:10.1126/science.abf2824 (2021).

52 Fujita, M. et al. The anisotropy1 D604N mutation in the Arabidopsis cellulose synthase1 catalytic domain reduces cell wall crystallinity and the velocity of cellulose synthase complexes. Plant Physiol 162, 74–85, doi:10.1104/pp.112.211565 (2013).

53 Bethke, G. et al. Pectin Biosynthesis Is Critical for Cell Wall Integrity and Immunity in Arabidopsis thaliana. Plant Cell 28, 537–556, doi:10.1105/tpc.15.00404 (2016).

54 Hibara, K.-I. et al. Abnormal Shoot in Youth, a homolog of molybdate transporter gene, regulates early shoot development in rice. American Journal of Plant Sciences 4, 1–9 (2013).

55 Benedetti, M. et al. Plant immunity triggered by engineered in vivo release of oligogalacturonides, damage-associated molecular patterns. Proc Natl Acad Sci U S A 112, 5533–5538, doi:10.1073/pnas.1504154112 (2015).

56 Pontiggia, D., Benedetti, M., Costantini, S., De Lorenzo, G. & Cervone, F. Dampening the DAMPs: How Plants Maintain the Homeostasis of Cell Wall Molecular Patterns and Avoid Hyper-Immunity. Front Plant Sci 11, 613259, doi:10.3389/fpls.2020.613259 (2020).

57 Cabrera, J. C., Boland, A., Messiaen, J., Cambier, P. & Van Cutsem, P. Egg box conformation of oligogalacturonides: the time-dependent stabilization of the elicitor-active conformation increases its biological activity. Glycobiology 18, 473–482, doi:10.1093/glycob/cwn027 (2008).

58 Rocha, P. S. et al. The Arabidopsis HOMOLOGY-DEPENDENT GENE SILENCING1 gene codes for an S-adenosyl-L-homocysteine hydrolase required for DNA methylation-dependent gene silencing. Plant Cell 17, 404–417, doi:10.1105/tpc.104.028332 (2005).

59 Moffatt, B. A. & Weretilnyk, E. A. Sustaining S-adenosyl-L-methionine-dependent methyltransferase activity in plant cells. Physiologia Plantarum 113, 435–442 (2001).

60 Patron, N. J. et al. Standards for plant synthetic biology: a common syntax for exchange of DNA parts. New Phytologist 208, 13–19 (2015).

61 Shimada, T. L., Shimada, T. & Hara-Nishimura, I. A rapid and non-destructive screenable marker, FAST, for identifying transformed seeds of Arabidopsis thaliana. The Plant Journal 61, 519–528 (2010).

62 Nelson, B. K., Cai, X. & Nebenführ, A. A multicolored set of in vivo organelle markers for co-localization studies in Arabidopsis and other plants. The Plant Journal 51, 1126–1136 (2007).

63 Clough, S. J. & Bent, A. F. Floral dip: a simplified method for Agrobacterium-mediated transformation of Arabidopsis thaliana. Plant Journal 16, 735–743, doi:10.1046/j.1365-313x.1998.00343.x (1998).

64 Schindelin, J. et al. Fiji: an open-source platform for biological-image analysis. Nature methods 9, 676–682 (2012).

65 Yang, W. et al. Regulation of meristem morphogenesis by cell wall synthases in Arabidopsis. Current Biology 26, 1404–1415 (2016).

66 Wightman, R., Wallis, S. & Aston, P. Hydathode pit development in the alpine plant Saxifraga cochlearis. Flora 233, 99–108 (2017).

67 de Reuille, P. B. et al. MorphoGraphX: A platform for quantifying morphogenesis in 4D. Elife 4, e05864 (2015).

68 Gilbert, H. J., Hazlewood, G. P., Laurie, J. I., Orpin, C. G. & Xue, G. P. Homologous catalytic domains in a rumen fungal xylanase: evidence for gene duplication and prokaryotic origin. Mol Microbiol 6, 2065–2072, doi:10.1111/j.1365-2958.1992.tb01379.x (1992).

69 Anthon, G. E. & Barrett, D. M. Comparison of three colorimetric reagents in the determination of methanol with alcohol oxidase. Application to the assay of pectin methylesterase. Journal of agricultural and food chemistry 52, 3749–3753 (2004).

70 Simmons, T. J. et al. Folding of xylan onto cellulose fibrils in plant cell walls revealed by solid-state NMR. Nat. Commun. 7, 13902 (2016).

71 Munowitz, M. G., Griffin, R. G., Bodenhausen, G. & Huang, T. H. Two-Dimensional Rotational Spin-Echo Nuclear Magnetic-Resonance in Solids - Correlation of Chemical-Shift and Dipolar Interactions. J. Am. Chem. Soc. 103, 2529–2533, doi:Doi 10.1021/Ja00400a007 (1981).

72 Hong, M. et al. Coupling amplification in 2D MAS NMR and its application to torsion angle determination in peptides. J. Magn. Reson. 129, 85–92, doi:DOI 10.1006/jmre.1997.1242 (1997).

73 Bielecki, A., Kolbert, A. C. & Levitt, M. H. Frequency-Switched Pulse Sequences - Homonuclear Decoupling and Dilute Spin NMR in Solids. Chem. Phys. Lett. 155, 341–346, doi:Doi 10.1016/0009-2614(89)87166-0 (1989).

74 Rienstra, C. M. et al. De novo determination of peptide structure with solid-state magic-angle spinning NMR spectroscopy. Proc. Natl. Acad. Sci. USA 99, 10260–10265, doi:10.1073/pnas.152346599 (2002).

75 Tsirigos, K. D., Peters, C., Shu, N., Käll, L. & Elofsson, A. The TOPCONS web server for consensus prediction of membrane protein topology and signal peptides. Nucleic acids research 43, W401–W407 (2015).

