## Supplemental data for "Discovery of putative Golgi S-Adenosyl methionine transporters reveals the importance of plant cell wall polysaccharide methylation"

A

Identity (upper triangle) / Similarity (lower triangle)

| | CeSAMT1 | GoSAMT1 | GoSAMT2 | GoSAMT3 |
|---|---|---|---|---|
| CeSAMT1 | – | 45% | 43% | 44% |
| GoSAMT1 | 72% | – | 84% | 80% |
| GoSAMT2 | 72% | 96% | – | 83% |
| GoSAMT3 | 73% | 96% | 98% | – |

B

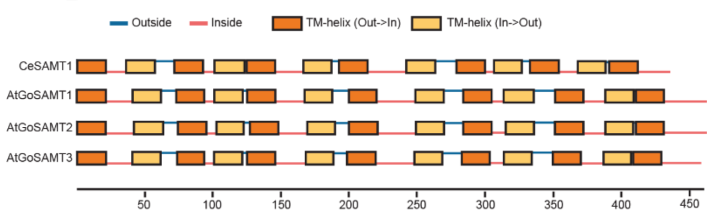


C

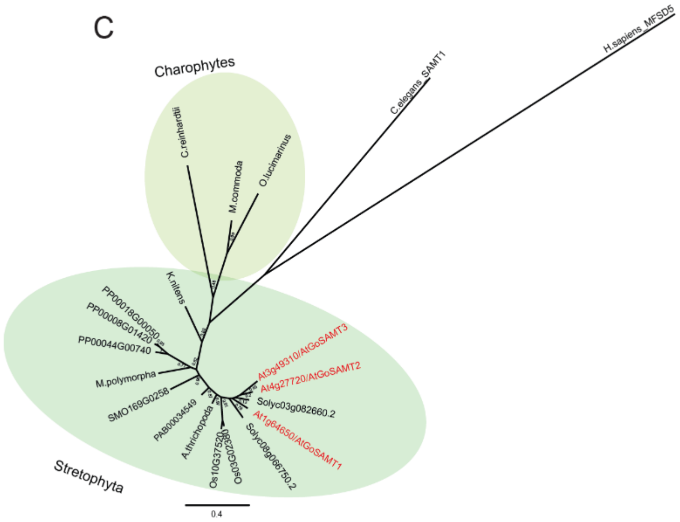


**Supplemental Figure 1**. **Arabidopsis GoSAMTs show similarity to CeSAMT1 and they are present throughout the plant kingdom. (A)** Sequence identity and similarity matrix generated based on MUSCLE alignments of whole length protein sequences of CeSAMT1 and Arabidopsis GoSAMTs. **(B)** CeSAMT1 and Arabidopsis GoSAMTs topology using TOPCONS [75]. **(C)** Phylogenetic tree of MFS_5 proteins from *Arabidopsis thaliana* (At), *Amborella thricopoda* (Atr), *Caenorhabditis elegans*, *Chlamydomona reinhardtii*, *Klebsormidium nitens*, *Homo* sapiens, *Micromonas commoda*, *Oryza sativa* (Os), *Ostreococcus* lucimarinus, Picea *abies* (PAB), *Physcomitrella patens* (Pp) *Selaginella moellendorffii* (SMO), *Solanum lycopersicum* (Solyc). Arabidopsis sequences are highlighted in red. Phylogenetic tree was generated using Molecular Evolutionary Genetics Analysis MEGA X and visualized by FigTree 1.4.2.

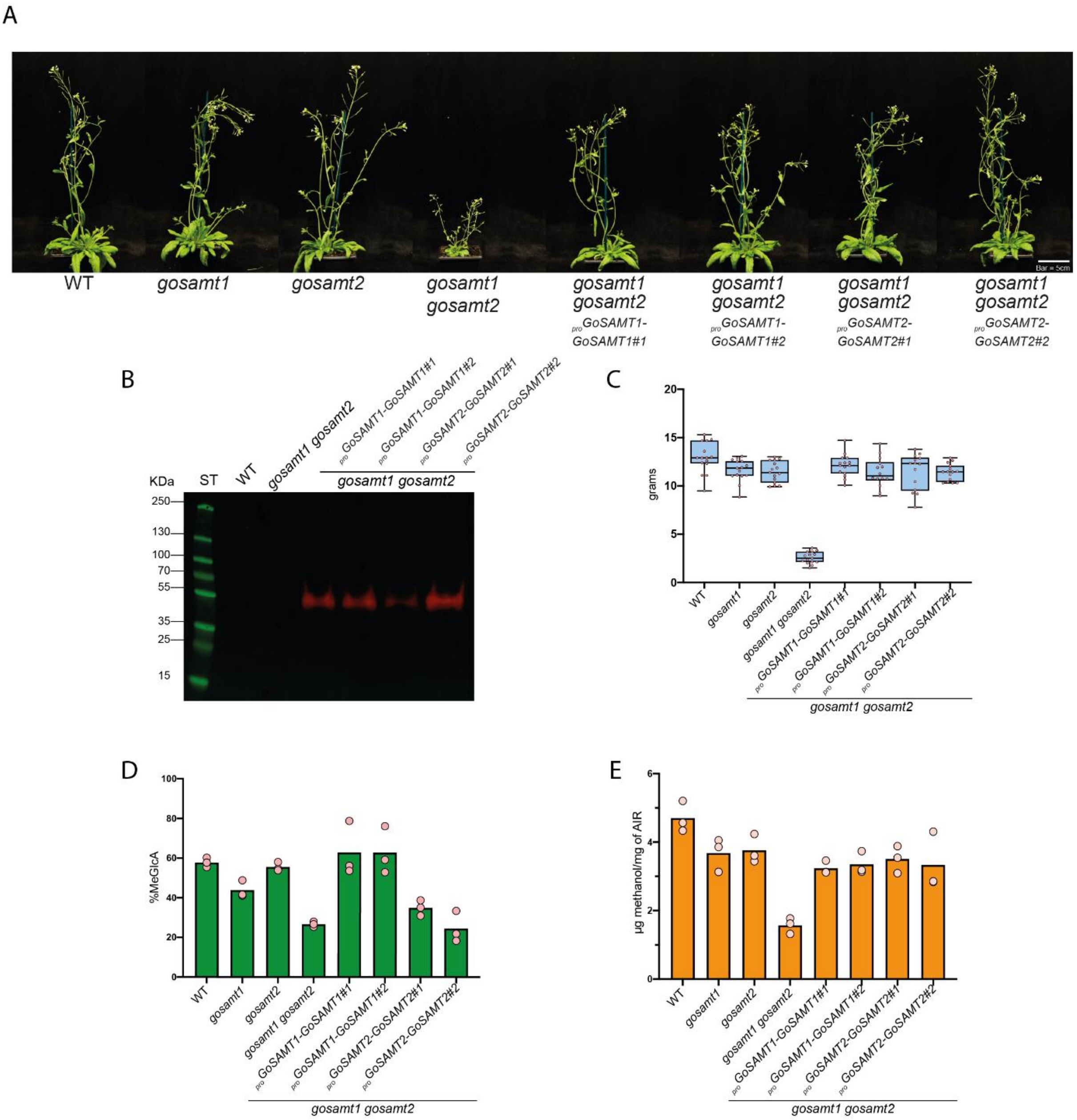


**Supplemental Figure 2. Complementation of *gosamt1 gosamt2* mutant. (A)** Pictures of adult plants of WT, *gosamt* mutants and *gosamt1 gosamt2* mutant molecular complemented lines. **(B)** Western blot, expression analysis of GoSAMT1-GFP and GoSAMT2-GFP complemented lines using anti-GFP. **(C)** Box and whiskers plot representing plant fresh weight of the different genotypes. **(D)** Ratio of released $Xyl_4GlcA$ and $Xyl_4{}^{Me}GlcA$ products after endoxylanase treatment of basal stem AIR, coupled to capillary electrophoresis experiments of secondary cell wall xylan of WT, *gosamt1*, *gosamt2*, *gosamt1 gosamt2* and molecular complemented lines. Values correspond to the mean of 3 biological replicates. **(D)** Methanol release experiments of leaf AIR of WT, *gosamt1*, *gosamt2*, *gosamt1 gosamt2* and molecular complemented lines. Values correspond to the mean of 3 biological replicates.

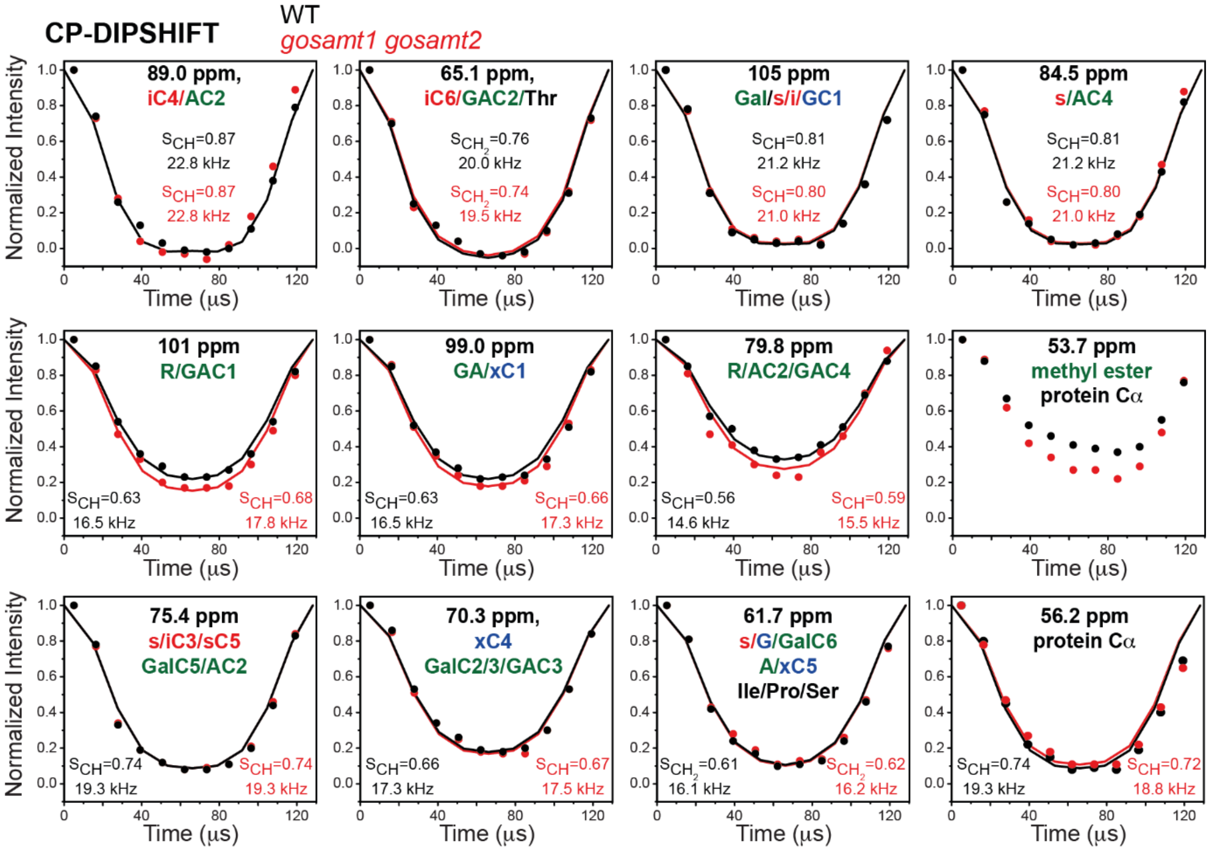


**Supplemental Figure 3. CP-DIPSHIFT experiment.** CP-DIPSHIFT curves of WT (black) and *gosamt1 gosamt2* (red) cell walls measured at 293 K. Best-fit $^{13}C$-$^{1}H$ dipolar coupling values (scaled by FSLG) and $S_{CH}$ order parameters are given in each panel. The experiment was conducted with C-H dipolar doubling, a CP contact time of 500 μs under 7.8 kHz MAS. The 101 ppm, 99.0 ppm, and 79.8 ppm peaks of pectin backbones are more rigid in the mutant than in the WT cell wall.

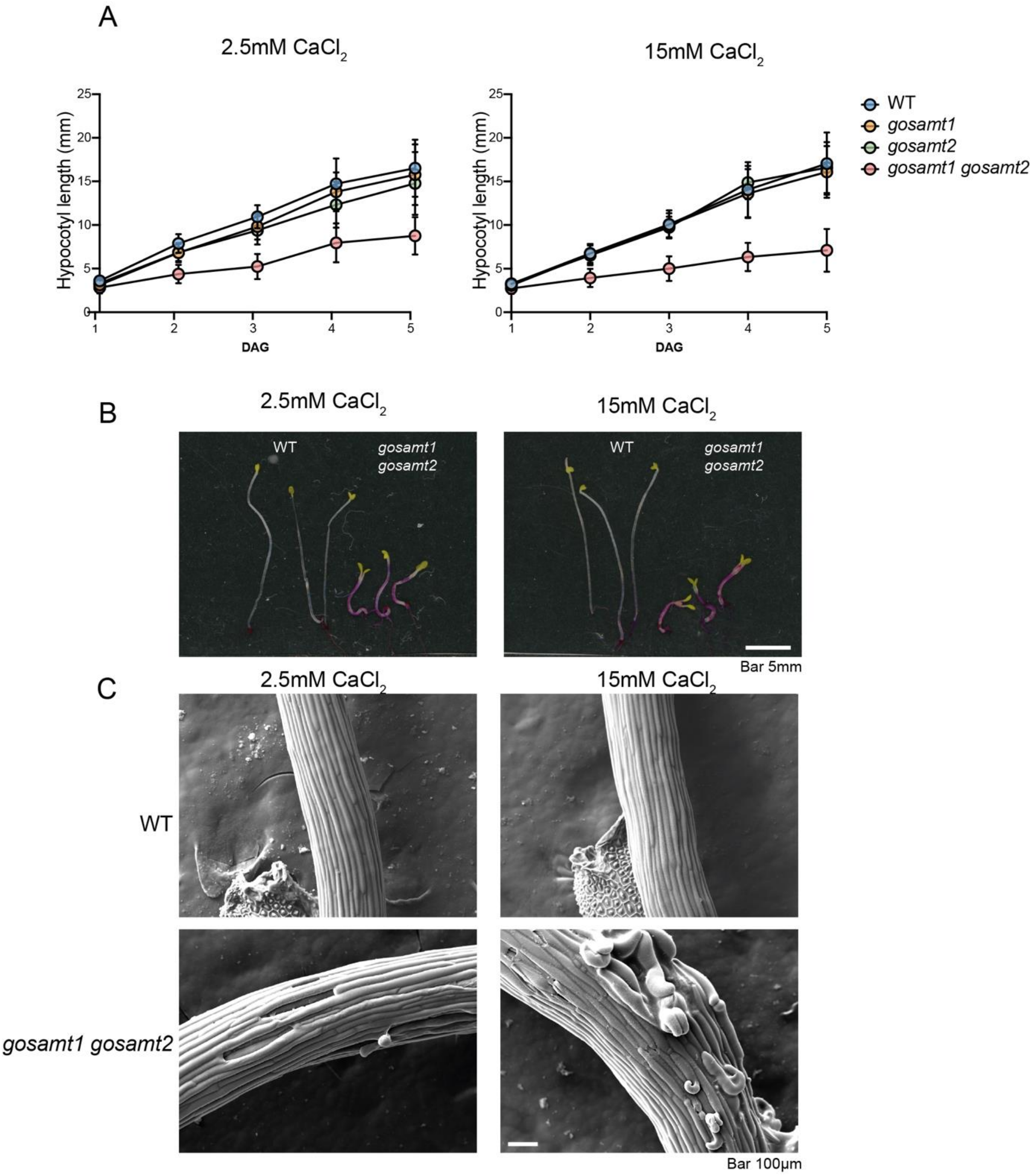


**Supplemental Figure 4. *gosamt1 gosamt2* display strong cell elongation and cell adhesion phenotypes in etiolated hypocotyls. (A)** Quantification of etiolated hypocotyls of WT, *gosamt1*, *gosamt2* and *gosamt1 gosamt2*, one to five days after germination (DAG) grown etiolated hypocotyls length grown in ½MS media and ½MS supplemented with $CaCl_2$ to a final concentration of 15mM. *n* ≥73 seedlings per genotype per time point. **(B)** WT and *gosamt1 gosamt2* hypocotyls etiolated hypocotyls length grown in ½MS media and ½MS supplemented with $CaCl_2$ to a final concentration of 15mM stained with ruthenium red. Bar 5mm **(C)** Scanning electron microscopy (SEM) of WT and in MS and in MS supplemented with $CaCl_2$ to a final concentration of 15 mM. Bar 100µm.

**Supplemental table 1. Primers used in this study**

| | |
|---|---|
| **Primers for Genotyping** | |
| *GoSAMT1* Left | TGTTCCAATTTGGAAATCGAG |
| *GoSAMT1* Right | CTTATGGCTTTGGAAAGGGAG |
| *GoSAMT2* Left | GAGATTCCTCCACCTTTCAC |
| *GoSAMT2* Right | TGGTGTTCTGTATCAGGGGTC |
| Salk Lb1.3 | ATTTTGCCGATTTCGGAAC |
| Gabikat o8760 | GGGCTACACTGAATTGGTAGCTC |
| **Primers for RT-PCR (5' to 3')** | |
| EF1α Forward | ATGCCCCAGGACATCGTGATTTCAT |
| EF1α Reverse | TTGGCGGCACCCTTAGCTGGATCA |
| GoSAMT1 Forward | TGGGTTTAGAGGGTCAGCTT |
| GoSAMT1 Reverse | ATGGAGATCTTCTACTTCGT |
| GoSAMT2 Forward | ATGGAGATTTTCTACTACTT |
| GoSAMT2 Reverse | TATGTTGAGGGGATCTTCTT |
| **Primers for construct sequencing** | |
| NOS terminator | TAATCATCGCAAGACCGGCA |
| 5’eGFP_reverse | GCTGAACTTGTGGCCGTTTA |
| Right border primer | AAACCTTTTCACGCCCTTTT |
| Left border primer | ATCGAGTGGTGATTTTGTGC |
